# The Central Role of the Tail in Switching Off Myosin II in Cells

**DOI:** 10.1101/606368

**Authors:** Shixin Yang, Kyoung Hwan Lee, John L. Woodhead, Osamu Sato, Mitsuo Ikebe, Roger Craig

## Abstract

Myosin II is a motor protein playing an essential role in cell motility. The molecule can exist as a polymer that pulls on actin to generate motion, or as an inactive monomer with a compact structure, in which its tail is folded and its two heads interact with each other. This conformation functions in cells as an energy-conserving storage and transport molecule. The mechanism of inhibition is not fully understood. We have carried out a 3D reconstruction of the switched-off form revealing for the first time multiple interactions between the tail and the two heads that trap ATP hydrolysis products, block actin binding, obstruct head phosphorylation, and prevent filament formation. Blocking these essential features of myosin function can explain the high degree of inhibition of the folded form of myosin, serving its energy-conserving, storage function in cells. The structure also suggests a mechanism for unfolding when activated by phosphorylation.

## Introduction

Myosin II is a motor protein that, together with actin filaments, generates mechanical force and motion using the chemical energy of ATP hydrolysis [4]. In muscle, myosin II is responsible for shortening and force production, while in nonmuscle cells it is essential in cell adhesion and division, intracellular transport and cell migration [5]. The myosin II molecule is a hexamer composed of two heavy chains, two essential light chains (ELC) and two regulatory light chains (RLC). The light chains and the N-terminal halves of the heavy chains form two globular heads, containing ATP- and actin-binding sites, while the heavy chain C-terminal halves form an α-helical coiled-coil tail extending from the heads. The tails self-associate to form the backbone of thick filaments, which are the functional form of myosin II in muscle and cell motility. Mutations in the heads and tail impair myosin function and cause muscle disease [6, 7]. Myosin II molecules can exist in two different conformations, 6S and 10S, named for their sedimentation coefficients [8]. 10S molecules have a compact structure [2, 9–12], in which the tail is folded into three segments of similar length, and the heads are bent back on the tail and interact with each other (Fig. 1). This folded conformation traps the products of ATP hydrolysis, causing strong inhibition of ATPase activity [13], and inhibiting assembly into thick filaments. Phosphorylation of the RLC in 10S molecules favors unfolding to the 6S form [10, 14, 15], in which ATPase activity is switched on [16] and the tail has an extended conformation, favoring filament assembly.

**Figure 1.**
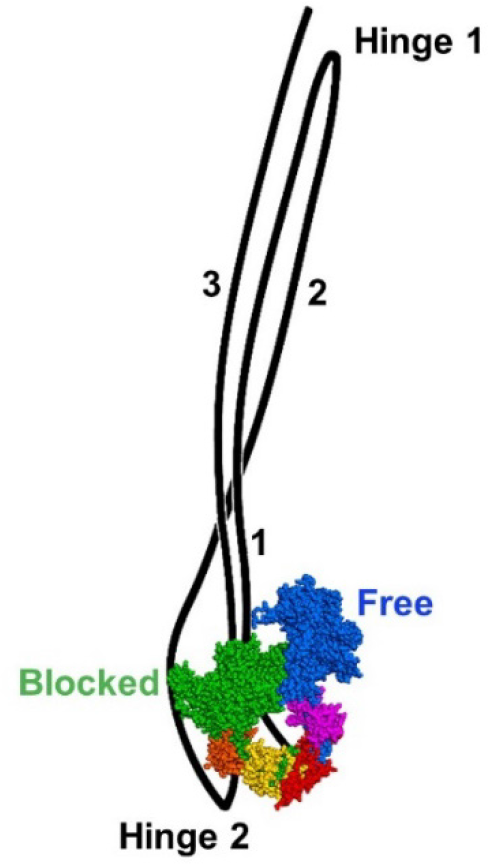
Model of 10S myosin. Blocked head consists of heavy chain (green), ELC (orange) and RLC (yellow). Free head shows heavy chain (blue), ELC (magenta) and RLC (red). The tail is folded into three segments: segment 1 (= subfragment 2), originating from the heads and ending at hinge 1; segment 2, starting at hinge 1 and ending at hinge 2; and segment 3, starting at hinge 2 and ending at the tip of the tail.

The inhibited, 10S molecule plays a fundamental role in nonmuscle cells, where it functions as an energy-conserving, storage form of myosin, and provides assembly units for thick filament formation when and where contractile activity is required [14]. Its compact conformation would facilitate rapid transport to the sites of filament assembly and thus motility. In smooth muscle, 10S myosin may also provide a pool of inactive monomers, which can unfold and augment myosin filament length or number when smooth muscle is activated [17–19]. In striated muscles, myosin is almost exclusively filamentous, but there is nevertheless a small concentration of 10S molecules, whose compact form has been proposed to facilitate transport to sites of filament assembly during muscle development and turnover of myosin filaments [20–22].

A unique structural feature of 10S myosin is an asymmetric interaction between its two heads (the interacting-heads motif, or IHM), which inhibits their activity [1]. Head-head interaction was first observed in two-dimensional (2D) crystals of smooth muscle heavy meromyosin in the switched-off state (RLCs dephosphorylated) [1], and later confirmed by electron microscopy (EM) and 2D classification of single myosin II molecules [9]. Inhibition was suggested to occur by different mechanisms in the two heads. In one head (“blocked”), actin-binding is hindered by proximity of the other head (“free”) to its actin-binding site. In contrast, ATPase activity of the free head is inhibited through stabilization of its converter domain, which interacts with the blocked head [1]. Three-dimensional (3D) reconstruction of thick filaments subsequently showed that 6S myosin, when polymerized into thick filaments, has a similar head-head interaction to that in folded, 10S molecules [23]. The IHM in thick filaments may underlie the super-relaxed state of muscle [24], in which myosin activity is highly inhibited, thus serving as an important energy-conserving mechanism for striated muscle.

The IHM structure has been compared in a variety of evolutionarily diverse species. EM and 2D image classification of negatively stained molecules shows that the compact, folded form of 10S myosin has been conserved since the origin of animals: head-head interactions in the IHM appear similar in all species studied [2, 25, 26]. Similarly, the IHM first observed in thick filaments in cryo-EM studies of tarantula muscle [23] has been confirmed in thick filaments from a variety of vertebrate and invertebrate species [27–30]. The high level of conservation of the IHM in isolated molecules and in thick filaments over hundreds of millions of years implies that it is a fundamental structure, critical to the function of muscle and nonmuscle cells [2, 25, 26].

While the inhibitory interactions between the heads in the IHM are similar in thick filaments and 10S molecules, there are likely to be critical additional interactions with the folded tail that account for the order of magnitude greater inhibition of 10S molecules compared with filaments [21, 31]. However, it is not known what these interactions might be, as no 3D reconstruction of the entire molecule has been reported. The interactions between the myosin heads in 10S myosin were well defined in studies of 2D crystals [32], but the tail was poorly visualized and the 3D reconstruction provided no insights into its potential role in regulation. The tail was clearly delineated in 2D class averages of isolated 10S molecules, but its course in three dimensions was not studied [9]. Here we present a 3D reconstruction of 10S myosin molecules using single particle analysis. The reconstruction clearly reveals the 3D course of the tail as it wraps around the heads, showing evidence for interaction with the heads at eight sites, which appear to inhibit actin-binding, ATPase activity, RLC phosphorylation and filament assembly by novel mechanisms. The results provide new insights into how the cell conserves energy, by a total shutdown of myosin II activity in its compact, storage form.

## Results

### 2D class averages of 10S myosin

Turkey gizzard smooth muscle myosin at physiological ionic strength (0.15 M NaAc) in the presence of ATP was observed by negative staining with 1% uranyl acetate, a technique providing key insights into protein domain structures and interactions [33, 34]. The compact, folded (10S) conformation dominated the EM images, with a small number of antiparallel 10S dimers, displaying a dumbbell-shaped structure (Fig. 2A) and very few extended (6S) molecules. Only 10S monomers were chosen for image processing. The tail in these molecules was folded into 3 segments, and wrapped around the heads, as previously described [9]. The heads showed different apparent shapes and sizes, corresponding to different orientations of the molecules on the EM grid (Figs. 2 B-E, S1). Compared to the relatively rigid arrangement of the interacting heads, the tail was quite flexible where it extended away from the heads. In this region, segments 1, 2 and 3 (Fig. 1) run closely together, and this combined 3-segment rod bends within a range of ~ 60° from its point of emergence from the heads, as previously described [9]. Typical 2D class averages of the interacting heads and the proximal portion of the 3-segment tail are shown in Figs. 2 B-E and S1. The asymmetric architecture of the blocked and free heads was consistent with the results of previous 2D classification analysis [2, 9, 25, 35]. Mirrored class averages, where molecules lie on the grid facing down or up, were observed as previously reported [35] (Fig. S1). The class averages showed that the myosin molecules typically lie parallel to the grid surface; the different orientations about their long axis enabled us to analyze the 3D structure of the molecules using single particle analysis.

**Figure 2.**
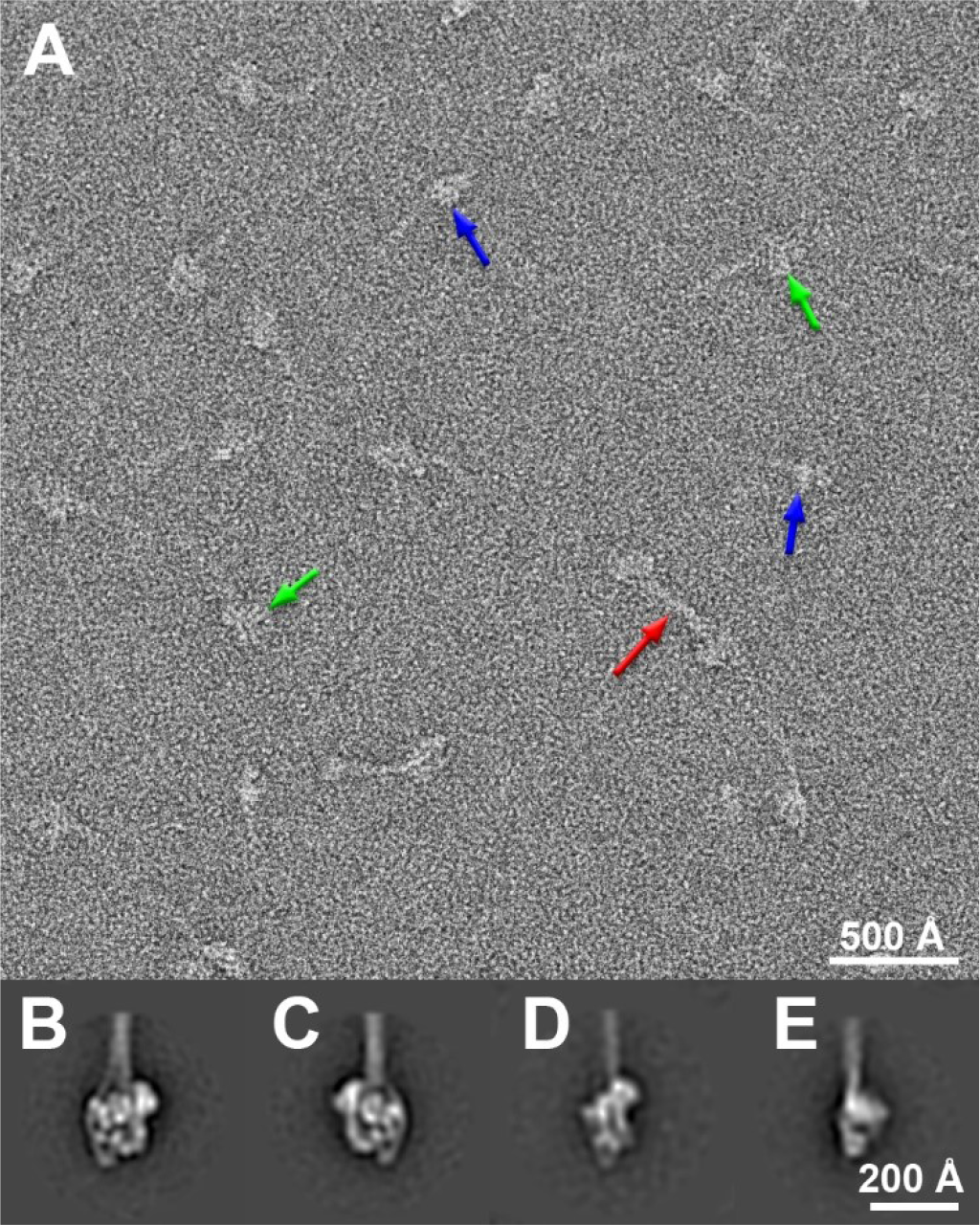
Negatively stained 10S smooth muscle myosin molecules. A. Two types of structure are visible: monomers, in front and side views (green and blue arrows, respectively), and occasional dimers (red arrow). B-D. 2D class averages of right and left views (B and C respectively [2]); (D), partially rotated view; and (E), side-view (see also Fig. S1).

### 3D reconstruction

An initial 3D structure of the folded molecule was computed using the random conical tilt method [36], in which images are recorded at 0° and 50° tilts. This model was then used for projection matching to determine the final single particle reconstruction (EMDB accession code EMD-20084). Computing the initial structure *ab initio* excludes potential model bias in the final reconstruction. The latter had a moderate resolution (25 Å; Fig. S2) due significantly to the flexibility of the molecule [9].

The head region of the reconstruction has the appearance of a flat disk, with an irregular polygonal “front” view (as viewed in Fig. 3A, D; Movie 1), enclosing a central hole, and a narrow side view (Fig. 3C, F, Movie 1), similar to the motifs seen in the 2D crystals of HMM and myosin [1, 32] and in thick filaments [23]. The atomic model of the interacting-heads structure computed from cryo-imaged smooth muscle HMM 2D crystals (PDB-1I84) [1] docked well into the reconstruction using rigid-body fitting (Fig. 4, Movie 2), confirming this similarity. This was clear in both front- and side-views (Fig. 4D, F). The fit implies excellent preservation of the IHM in negatively stained single molecules (*cf*. [33]), without significant flattening, and enables us to interpret head structure and interactions at higher resolution (<7 Å) than the reconstruction itself [37].

**Fig. 3.**
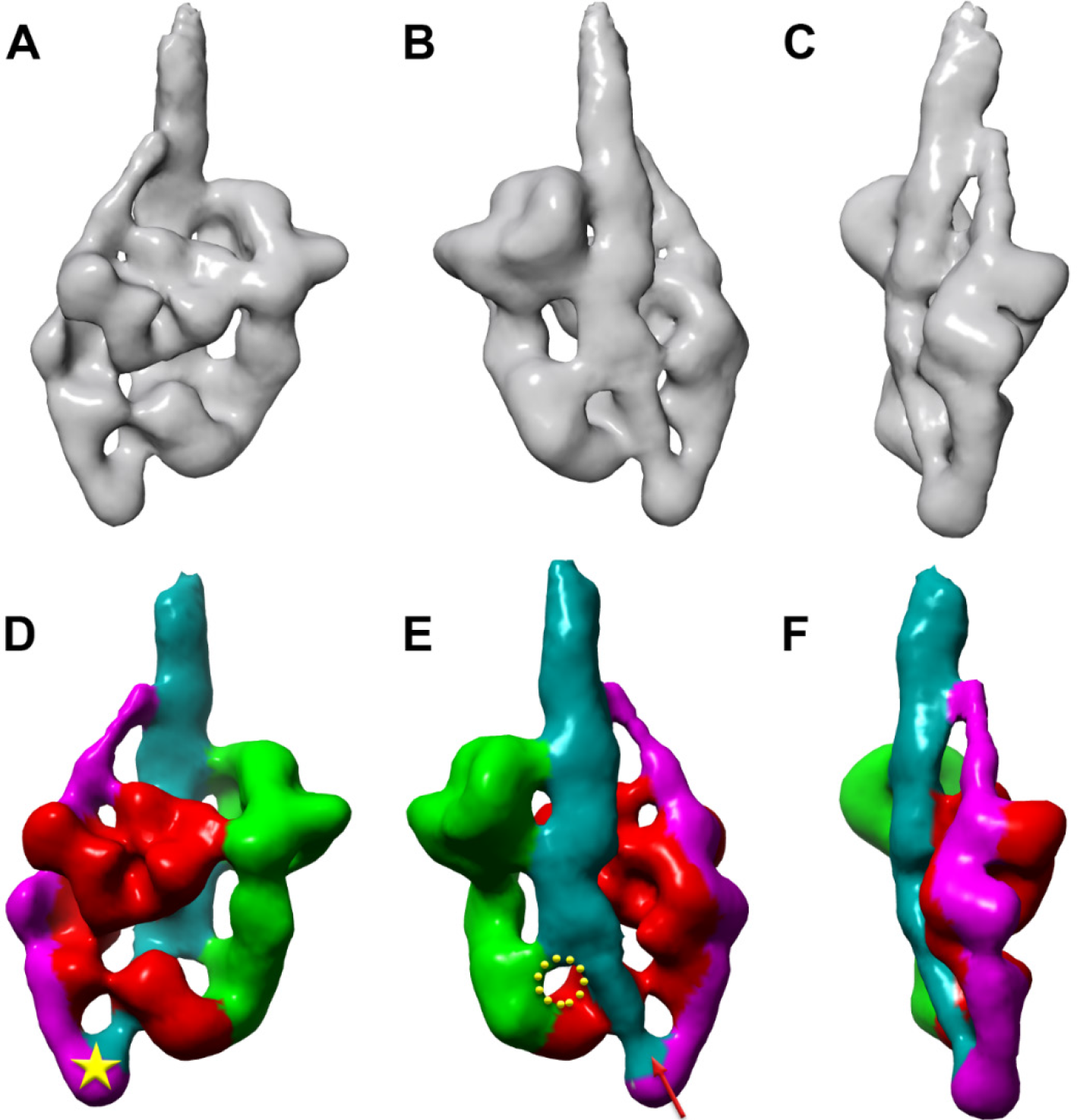
3D reconstruction of 10S smooth muscle myosin (EMDB accession code EMD-20084). (A), (B) and (C) are front, back and side views, respectively. (D), (E) and (F) correspond to (A), (B) and (C), with colors indicating different components of the structure. Blocked head is red; free head, green; segments 1 and 3, cyan; segment 2, magenta. In (E), yellow circle indicates putative location of missing segment 1 density (see Fig. S3); red arrow, start of segment 3. Yellow star in (D) indicates the second hinge point.

The reconstruction also reveals, for the first time, the course of the folded tail as it wraps around the heads – a feature that was mostly missing from the only previous reconstruction of 10S myosin [32]. Only a small portion of the folded tail (~150 Å) extending as three merged segments above the heads was included in the reconstruction, due to flexibility of the distal region. However, the essential part of the tail that interacts with the heads was included. Three segments of the tail could be identified in this region (Fig. 3). Strikingly, all three appear to bind to the heads: segment 2 along the left side of the blocked head (Figs. 1, 3A, D), and segments 1 and 3 on the back side of both heads (Fig. 3B, E, Movie 1). Segment 1 (subfragment 2 of the tail) would be expected to start at the merge point of the lever arms of the blocked and free heads (blue ribbon, Fig. S3). However, there was missing density for this initial portion of subfragment 2 in the reconstruction (circle, Figs. 3E, S3), a known issue for this very flexible region of the tail [28, 38]; more density becomes visible at low contour cutoff (Fig. S4). The first visible part of segment 1 density was assumed to occur where the tail density (cyan) widens, as segment 1 joins segment 3 (above the circle, Fig. 3E). Segment 1 then runs next to segment 3 across the blocked head (in a region now known as the mesa [39]) towards the top, where segments 1 and 3 merge with segment 2 (Fig. 3D, E). Segment 2 (magenta in Fig. 3) branches from the merged segments near the top and wraps around the blocked head. The end of this segment and the starting point of segment 3 form the second hinge point of the tail (Figs. 1, 3D), thought to occur at Glu1535 [9]. This region of density (marked with a star in Fig. 3D) is unique to 10S myosin, and not found in HMM (lacking segments 2 and 3) or thick filaments (where the tail is unfolded). Segment 3 (red arrow in Fig. 3E) starts from the hinge point, then runs, merged with segment 1, over the back of the blocked head (Fig. 3D, E) to the top of the IHM where they join segment 2. This arrangement of segments 1 and 3 on the same side of the IHM (Figs. 3, 5) would allow for rapid opening of the molecule upon activation. An alternative arrangement, consistent with 2D projection images (Figs. 2, S1; [9]), but not with the 3D structure, would have segments 1 and 3 on opposite sides, sandwiching the blocked head and restricting opening of the IHM (*cf*. [9]).

**Figure 4.**
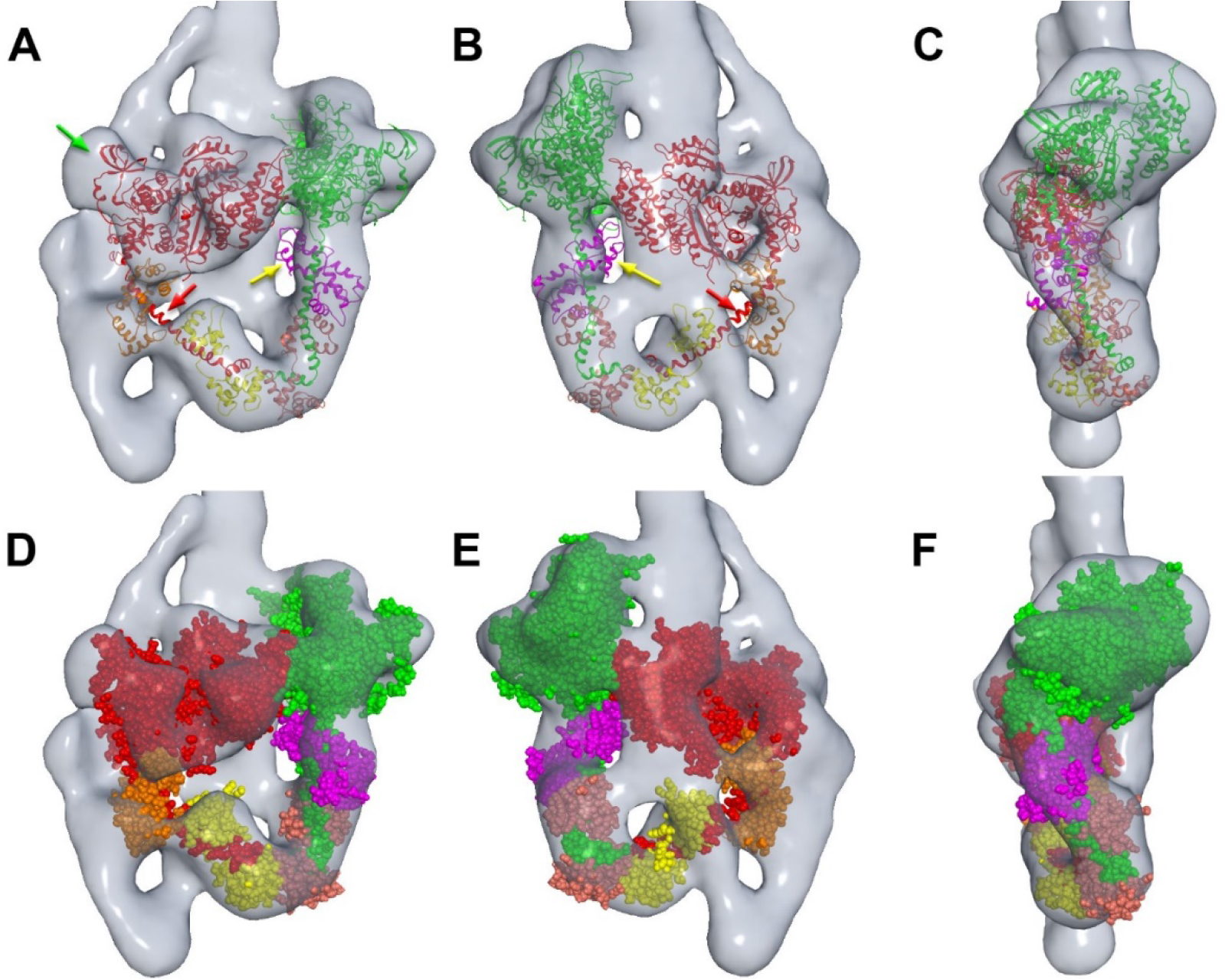
Docking of atomic model of the IHM into the reconstruction. The 2D crystal model of smooth muscle myosin (1i84; [1]) was used in the fitting. Molecular dynamics flexible fitting was not used, due to the low resolution. (A), (B) and (C) are front, back and side views, respectively, docked with ribbon representation of the atomic model. (D)-(F) correspond to (A)-(C), but with space-filling model. The atomic model fits well into the reconstruction in all views, except for the narrow part of the lever arm in the blocked head (red arrow) and part of the ELC in the free head (yellow arrow). Unfilled densities mostly represent portions of the three segments of the tail, for which there is no atomic model. There is also some unfilled density in the blocked head (green arrow) that would be filled by repositioning the flexibly-connected SH3 domain (not done with the rigid-body docking procedure; see Methods). In a previous study, molecular dynamics flexible fitting of cryo-imaged tarantula thick filaments demonstrated this point [3]. Note: (C) and (F) rotated in opposite direction from (C) and (F) in Fig. 3.

### Intramolecular interactions within 10S smooth muscle myosin molecules

The 3D reconstruction establishes key interactions occurring within the 10S myosin molecules (summarized in Table 1). Three types of interaction were found: head-head, head-tail, and tail-tail. The interaction between the blocked (B) and free (F) heads (BF) has been described before [1, 32], and is confirmed here. Eight putative interactions between the heads and the tail (T) have not previously been observed in 3D: these include the free head with tail segments 1 and 3 (TF), and the blocked head with segments 1, 2 and 3 (TB). Interactions between different segments of the tail were also observed (TT).

**Table 1:** Summary of IHM interactions.

| Interaction | Interacting components | Molecule/filament | Cartoon |
| --- | --- | --- | --- |
| BF | Blocked head, free head | 10S, filament |  |
| TB1 | Segments 1 and 2, blocked head mesa | 10S, filament |  |
| TB2 | Segment 2, blocked head SH3 | 10S, filament |  |
| TB3 | Segment 2, blocked head converter | 10S, filament |  |
| TB4 | Segment 2, blocked head ELC | 10S |  |
| TB5 | Segment 3, blocked head RLC | 10S |  |
| TF1 | Segment 1 or 3, free head actin-binding loop | 10S, filament |  |
| TF2 | Segment 1 or 3, free head ATPase loop 5 | 10S |  |
| TF3 | Segment 1 or 3, free head RLC | 10S |  |
| TT1 | Segments 1, 2, 3 | 10S |  |
| TT2 | Segments 1, 3 | 10S |  |

#### Head-head interactions

As described previously, the overall conformation of the IHM is produced by the blocked and free heads (red and green respectively in Fig. 3) interacting with each other (BF, blue arrow in Fig. 5A). The atomic model from this earlier work (PDB1i84) can be fitted well into the volume (Fig. 4, Movie 2).

#### Tail-head interactions

Three interactions are found between segment 2 of the tail and the blocked head as the tail wraps around its perimeter (Fig. 5A). TB2 is the interaction between tail segment 2 and the blocked head SH3 domain (blue in Fig. 5A). The blocked head is likely to be stabilized by this interaction. Interaction TB3 was found between segment 2 and the converter domain of the blocked head (cyan in Fig. 5A). The third interaction, TB4, was found between segment 2 and the ELC of the blocked head (orange in Fig. 5A).

**Figure 5.**
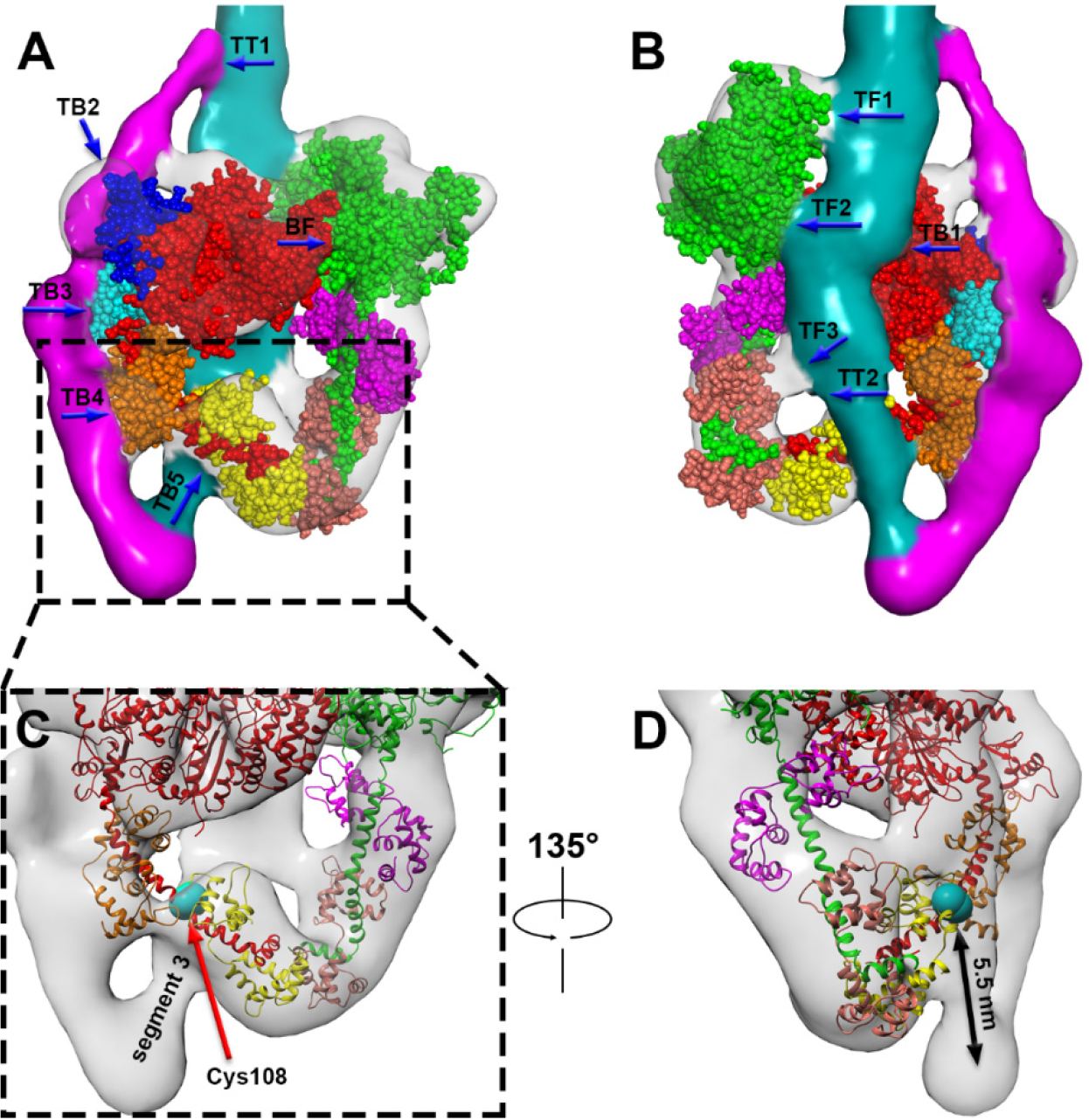
Interactions between the tail and heads in 10S myosin. (A) Front view shows the interactions between segments 2 (magenta) and 3 (cyan) and the blocked head (TB2, TB3, TB4, TB5), between the two heads (BF), and between segment 2 and the other tail segments (the interaction extends upwards from TT1). TB2 is between the SH3 domain (blue) in the blocked head and segment 2, which lies under it. TB3 is between segment 2 and the converter domain (cyan) of the blocked head. TB4 is between segment 2 and the ELC (orange) of the blocked head. TB5 is between segment 3 and the blocked head RLC. (B) Back view shows the interaction between the free head and the tail (TF1, TF2, TF3), between segments 1 and 3 (unresolved from each other) and the blocked head (TB1), and between segments 1 and 3 (starting at TT2). Interaction TF1 is between the actin-binding loop in the free head and the tail, while TF2 is between the free head and segment 1 or 3. TF3 may represent merging of tail and blocked head densities and not be a real interaction (see text). TB1 is an extended interaction between the blocked head and segment 1 and/or segment 3. (C) Magnified view of TB5. Cys108 is represented by a cyan sphere. (D) The view in (C) is rotated 135° to visualize the geometric relationship between Cys108 and segment 3.

Four interactions were observed between the tail and the underside of the heads (Figs. 5A, B, Table 1). TB1 is an extensive interaction, where the tail (including segments 1 and 3, which are not resolved from each other) runs over the back-surface of the blocked head motor domain (Fig. 5B). TB5 involves interaction between the RLC of the blocked head and segment 3, close to the second tail hinge point (Fig. 5A). TF1 occurs between the tail and the actin-binding loop of the free head, while TF2 is the interaction between tail segment 1 or 3 and the upper 50K domain of the free head. TF3 is a possible interaction between the tail and the free head RLC, but could alternatively represent low resolution merging of densities without actual contact (Fig. S4; see Discussion). These interactions between the tail and the free head were not reported in 2D analysis of smooth muscle myosin [9, 35], probably because the tail in these interactions is superimposed on the heads and cannot be identified in the 2D images.

#### Tail-Tail Interactions

In addition to the head-tail interactions, two regions of contact were found between the different segments of the tail. TT1 is an extensive interaction starting at the top of the IHM (Fig. 5A), where segment 2 merges with segments 1 and 3, and continuing up as a triple coiled-coil complex [9]. A second extensive interaction appears likely between segments 1 and 3 (unresolved from each other), as they pass together over the blocked head (Fig. 5B). This interaction starts at TT2, where the first part of segment 3 (cyan in Fig. 5B), would merge with the initial portion of S2 (not visible in the reconstruction, as discussed earlier). Other parts of the tail (the region of segment 2 peeling away from the triple segment and wrapping around the blocked head, and the start of segment 3 after the second tail hinge) exist as single coiled-coils, consistent with their narrower diameter in the reconstruction.

### Heterogeneity in the reconstruction

The flexibility of the merged tail segments extending beyond the head region is visualized in raw images, and was investigated using 2D classification by Burgess *et al.* [9], who showed that the tail flexed within a range of ~ 60° where it left the heads. The heads are also flexible, though more rigid than the tail. 3D classification was performed to explore flexibility and heterogeneity in the 10S molecules. 15833 particles were classified into 6 classes using RELION [40] (Figs. 6, S5). Classes 3 and 4, comprising 51% of the particles, exhibited very similar structures (Fig. 6C, D). Both tail and IHM were visualized clearly in 3D reconstructions and could be fitted well with the 3D atomic model of chicken smooth muscle myosin. The particles in these classes were combined and used for 3D refinement to produce the reconstruction described above. The other 4 classes were smaller and exhibited different tail and IHM structures in their reconstructions (Fig. 6A, B, E).

The classification results suggest that the tail plays a crucial role in stabilizing the compact structure of 10S myosin. When the tail can be traced explicitly (classes 3, 4 and 6), the IHM shows more consistent structural features than in the other three classes. 30% of myosin molecules (classes 1, 2 and 5) had flexible or disordered tails. Segment 2 density was discontinuous in class 1 and almost completely disappeared in classes 2 and 5, and the second hinge point was not visible in the latter two (Fig. 6B, E). Although these classes exhibited a similar basic asymmetric organization of heads, their IHM structure was somewhat varied, and the atomic model of the IHM (PDB 1I84) could not be docked into them very well. This implies that the tail directly influences the stability of the molecules, helping to maintain a compact structure: the weak interaction between the blocked and free heads is strengthened by the interactions of the heads with the tail. Disruption of these interactions results in conformational changes and less compact myosin molecules. Class 6 molecules exhibited less detail than classes 3 and 4, and were therefore also excluded from the final reconstruction.

**Figure 6.**
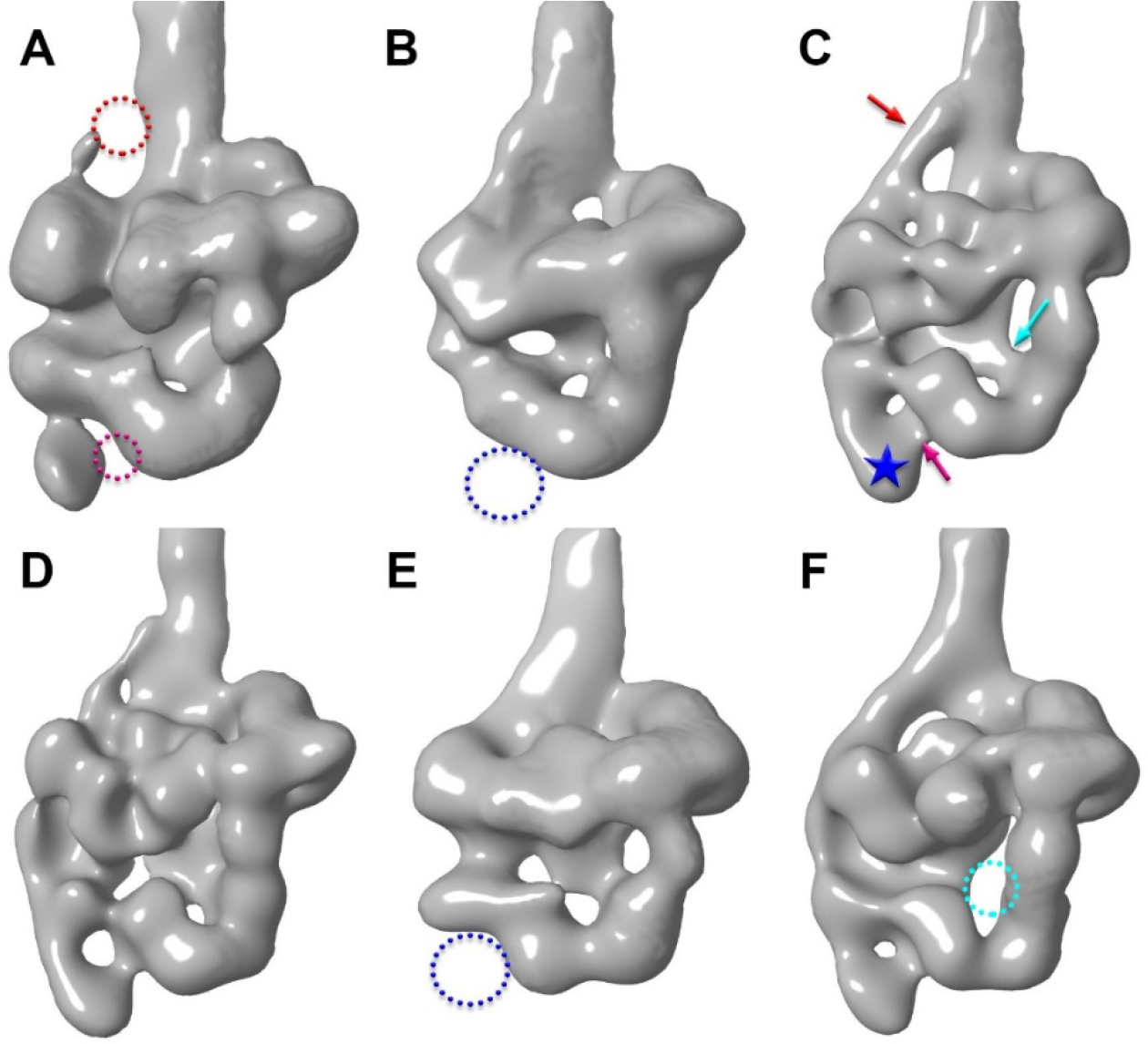
Heterogeneity in the reconstruction. 3D classification using RELION produced six classes of conformation (A-F show classes 1-6 respectively), showing that the 10S myosin molecules are quite heterogeneous.(A) The tail density partially disappeared in some regions (red dotted circles, *cf*. red arrows in (C), where it is visible). (B) The tail cannot be explicitly identified in this reconstruction and the second hinge point (star in (C)) is missing (blue circle). The reconstructions in (C) and (D) showed similar structural features, with both the interacting heads and the tail clearly visible. (E) Segment 2 around the blocked head is not visible, and the second hinge point is absent (blue circle), as in (B). (F) The tail wraps around the blocked head, but the density present in (C) (cyan arrow) and (D)
is absent (cyan circle).

## Discussion

While the head interactions that define the IHM in 10S myosin have previously been studied in detail [1, 32], the role of the tail in inhibiting function has not been well appreciated. Our reconstruction shows the 3D organization of the tail for the first time, providing crucial insights into the mechanism of inhibition that underlies the storage function of the folded molecule. The densities of the tail running across and wrapping around the two heads can be traced explicitly (Fig. 3D, E, F, Movie 2), in contrast to earlier studies where the tail could not be identified [1] or was not analyzed in 3D [9]. These earlier studies illustrate the difficulties of visualizing the myosin tail by cryo-EM and the ease with which it can be seen by negative staining, our technique of choice (see [33, 34] and Methods). While the resolution of the reconstruction is not high (due partly to flexibility of the molecule), we could easily detect the different domains of the myosin heavy chain and light chains, revealing steric blocking of specific regions of the heads by the tail that explains how the molecule is so effectively switched off in the quiescent state [31]. Fitting the atomic model of the IHM (PDB 1i84) to the reconstruction further refined our interpretation of interactions [37][]. The most important finding from our study is that interactions of the tail with the heads and with itself, combined with interactions between the heads, block all four key functional characteristics of 10S myosin: 1. ATPase activity, by head-head and head-tail interaction; 2. Actin binding, by head-head and head-tail interaction; 3. Activation by phosphorylation, through RLC-tail interaction; and 4. Filament formation, by tail-tail interaction. Inhibition of every aspect of myosin function appears to be nature’s failsafe method for conserving energy in the inactive state.

### The tail plays a central role in switching off 10S myosin activity

The reconstruction shows evidence for eight possible interactions between the tail and the heads in 10S myosin (Table 1). Five of these involve the blocked and three the free head, suggesting that the blocked head is the more stable [9]. All eight interactions may be important in creating a fully inhibited 10S molecule, but two stand out as being the most stable.

#### 1. Segment 1-blocked head interaction

Interaction between the blocked head and segment 1 (subfragment 2; TB1; Fig. 5B) appears to be one of the most fundamental interactions of the inhibited state, helping to maintain the blocked head in its folded-back conformation. This is suggested by its occurrence not only in myosin filaments and 10S myosin, but also in HMM [9]. In HMM, the free head preferentially detaches from the IHM in some molecules, while the blocked head remains bound to subfragment 2 [9]. The free head also detaches more readily than the blocked head in full-length (10S) myosin [9] (Fig. S6) and filaments [41], further suggesting stabilization of the blocked head by its binding to subfragment 2. Smooth muscle HMM constructs with different tail lengths are switched off only when the tail is equal to or greater in length than the head [42]. This is approximately the distance segment 1 must travel from its origin at the head-head junction to its interaction (TB1) with the blocked head mesa (Fig. 5B). This supports the view that the S2-blocked head interaction plays a critical role in inhibition [23].

#### 2. Segment 3-blocked head RLC interaction

We suggest that interaction TB5 (Fig. 5A), between segment 3 and the blocked-head RLC, is the strongest (and therefore the key) interaction that pins the tail to the heads, creating the folded, inhibited structure of 10S myosin, and helping to hold segments 2 and 3 in position where they interact with key sites on the blocked and free heads. The existence of this interaction is supported by photo cross-linking, showing that Cys108, in the C-terminal half of the RLC, is cross-linked to the tail between Leu1554 and Glu1583, which lie in segment 3 [35]. The tail must therefore fold back on to the light chain domain. The calculated distance from Glu1535 to Lys1568 (the middle residue in the cross-linking region) is ~ 48 Å, assuming 1.485 Å rise per residue along a coiled-coil. In the reconstruction, the distance measured from Glu1535 (hinge 2 [9], star in Fig. 3D) to Cys108 (cyan in Fig. 5C, D; Movie 3) in the blocked head RLC is similar (~ 55 Å). In the crosslinking studies, only one of the two RLCs is crosslinked to the tail [35, 43]. This is readily explained by the reconstruction, which shows that only blocked head Cys108 is close enough to the tail for crosslinking (Movie 3).

The crucial importance of TB5 to the folded structure is illustrated by previous biochemical and EM observations of 10S myosin. **(1)** When observed by rotary shadowing or negative staining, molecules that are not crosslinked by glutaraldehyde show a loosely folded structure, in which the only (and therefore strongest) intramolecular contact occurs between the start of segment 3 and the neck region (containing the RLC) of one head [10, 35]: the other interactions observed in our work are presumably weaker, and, in the absence of the glutaraldehyde treatment that we used, are disrupted by binding of the molecule to the negatively charged mica surface used for shadowing [26]. **(2)** In *Dictyostelium* and many insect flight muscle myosin II molecules, segment 3 does not bind to the blocked head; in these molecules, the tail and free head are quite flexible (Fig. S6A, C). **(3)** Segments 1 and 2 of smooth muscle myosin are together 1020 Å long [9]. Myosin II’s from amoeba (*Acanthamoeba castellanii*) and yeast (*Schizosaccharomyces pombe*) are much shorter than smooth muscle myosin, with <u>total</u> tail lengths of 870 and 910 Å, respectively [26]. They thus lack segment 3 entirely and the last 100-150 Å of segment 2. Neither of these myosins folds [26], further supporting the idea that segment 3 is essential for forming the 10S structure. **(4)** Biochemical observations also demonstrate the importance of the RLC for folding. The folded conformation fails to form when the RLC is removed [44] or replaced by skeletal RLC [45]. Experiments with chimeric smooth/skeletal RLCs show that native smooth N- and C-terminal lobes are both required for formation of 10S myosin [45]—the C-terminal lobe for full binding of the RLC to the heavy chain, and the N-terminal for folding. TB5 in our reconstruction shows segment 3 positioned over the C-terminal lobe and immediately next to the N-terminal lobe of the blocked head RLC, providing a structural explanation for these findings (Fig. 7E, F). The first twenty-four residues of the RLC are absent from the N-terminus in PDB 1i84, probably due to mobility. This region (dubbed the phosphorylation domain (PD) [46]) stabilizes tail folding in smooth muscle myosin [47]. We have estimated the location of these residues by superimposing the RLC of scallop (PDB 3JTD, which includes eleven additional N-terminal residues) on to the smooth muscle structure (Fig. S7). Residues 1-11 in scallop (14-24 in smooth muscle) of the blocked head RLC appear to contact segment 3, consistent with this stabilizing interaction. Based on sequence analysis and modeling, it has been suggested that the N-terminal 24 amino acids, including a specific group of positively charged residues, may lie over a cluster of negatively charged residues in segment 3, trapping this region of the tail on top of the RLC C-lobe and accounting for the strength of this interaction [35]. An alternative proposal is that these N-terminal RLC residues interact with helix A of the ELC in the switched off molecule [48].

#### 3. Other interactions

While the blocked head RLC interacts with segment 3, a putative interaction (TF3) with the free head RLC is less certain (Fig. 5C, D). The density in this region is very weak and fitting of PDB 1i84 suggests that there is no RLC density to fill the volume (Fig. S4); the relatively uniform cylindrical tail also lacks such density (Figs. 4, 5, S3, S4). This apparent interaction could therefore represent low resolution merging of the tail and head densities without actual contact. TF3 is close to the point where the first portion of segment 1, missing from the reconstruction, as discussed earlier (Figs. 3E, S4), would meet segment 3. This may account for the sudden widening of the tail at this location (Fig. 5B), leading to possible fusion with the RLC density.

In addition to interacting with the heads, the folded tail appears to interact with itself, where segments 1 and 3 run next to each other over the blocked head (interaction starting at TT2; Fig. 5B), and at the top of the reconstruction, where all three segments form a compact, rod-shaped structure, extending upwards from TT1 (Figs. 3, 5A). These interactions appear to play an essential role in the function of 10S myosin. First, folding sequesters the tail, inhibiting polymerization with other tails to form filaments, the active form of myosin II in cells. Second, extending up from the TT1 contact site, the complex of the three tail segments is quite flexible and can be significantly curved [9]. However, the interacting-heads structure remains relatively rigid. The TT1 contact site may function as a lock, preventing conformational changes in the curved rod complex from being transferred to the heads, thus helping to maintain their inhibited state. This is suggested by our observation that in the absence of TT1, as in *Dictyostelium* (Fig. S6B) and many insect indirect flight (Fig. S6D) muscle myosin molecules [26], the free head frequently detaches from the blocked head. The interaction between segments 1 and 3 starts at TT2. Proximity of TT2 to interaction TB5 (Fig. 5) may stabilize the interaction between segment 3 and the RLC in the blocked head.

### Tail segments 2 and 3 stabilize the IHM in 10S myosin

The atomic model of the IHM (PDB 1i84, based on 2D crystals of HMM [1]) fits well into the head densities of the reconstruction, without the need for any substantial change in head structure (Fig. 4D-F, Movie 2). This shows that binding of tail segments 2 and 3 (absent from HMM) to the heads in 10S myosin is not required to generate the head-head interactions, and that it does not significantly distort them. However, the frequency of formation of the interacting-heads structure in HMM, containing only the first third of the tail and having only one interaction between the tail and heads (TB1 in Fig. 5B), and the stability of the free head in HMM, are both lower than in full-length (10S) myosin [9]. Thus, these segments stabilize the head-head interactions, the compact structure and the inhibited function of 10S myosin. Several observations support this view: **(1)** a subset of the reconstructions that lack some of the tail density features correspondingly had a more varied head structure (Fig. 6A, B, E). **(2)** Myosin from *Dictyostelium* [26], which lacks segment 2 wrapping around the blocked head [26], showed more frequent cases of non-interacting heads (Fig. S6C, D). **(3)** Myosin molecules from insect flight muscle frequently behaved similarly (Fig. S6A, B). These observations suggest that the interactions of all three tail segments with the heads are crucial in fully stabilizing the structure of 10S myosin.

### Intramolecular interactions in thick filaments and single molecules: similarities and differences

10S myosin and thick filaments share several interactions. These include BF, TB1, TB2, TB3 and TF1 (Fig. 5, Table 1). Conservation of these interactions between filament and molecule suggests that they play an important role in the inhibited state of myosin. In both filament and molecule, it is thought that the BF interaction, between the two heads, constrains movement of the converter domain of the free head, inhibiting phosphate release, and blocks actin-binding by the blocked head [1, 23]. TB1 occurs between segment 1 and/or segment 3 and the blocked head where these two segments run over its motor domain (Fig. 5B). In filaments, where the tail is not folded [23], and in HMM, where segments 2 and 3 are absent [9, 35], this blocked head-S2 interaction is also present, supporting the view that TB1 involves S2, and consistent with data showing that subfragment 2 binds to subfragment 1 in solution [49]. The reconstruction suggests that in 10S myosin, segment 3 also binds to the blocked head motor domain. The tail is wider where it runs over the blocked head than at the start of segment 3, where it is a single α-helix (Fig. 5B), suggesting that segment 3 runs next to segment 1 over the blocked head. This double interaction with the blocked head would presumably help to stabilize the IHM of the folded molecule. TB2 (Fig. 5A) occurs between segment 2 and the SH3 domain (blue in Fig. 5A) of the blocked head. A related interaction is observed in relaxed thick filaments, between the SH3 domain of the blocked head and the tail from the next pair of heads away from the bare zone [23, 50]; however, in this case the interaction involves segment 1, and its polarity is the reverse of that in the 10S molecule. TB3 occurs between segment 2 and the converter domain of the blocked head (Fig. 5A). As with TB2, a similar interaction (but with reverse polarity) occurs between the converter domain and segment 1 of the neighboring tail in thick filaments [23]. The fifth interaction, TF1, occurs between the tail and the free head actin-binding loop in 10S myosin. A similar interaction between S2 and the actin-binding loop has been suggested in tarantula thick filaments [51], suggesting that TF1 also involves S2.

The other interactions in 10S myosin are unique to the folded structure. These tail-head and tail-tail interactions are presumably important specifically to the switched-off, folded structure, which underlies its storage and transport function. The compact structure would make transport more efficient through the crowded intracellular environment, while inhibition of ATPase and actin binding would minimize energy use and futile binding to actin filaments during transport.

### 10S myosin is switched off by steric blocking of ATPase and actin-binding sites on both heads

EM studies of 10S smooth muscle myosin [1, 32] suggested that inhibition was achieved by different mechanisms in the two heads: (1) by preventing actin binding in the blocked head; and (2) by inhibiting ATPase activity in the free head [1]. But they provided no information on the possible role of the tail in inhibition. Our results suggest that the tail plays a central role in switching off activity, such that *both* of these inhibitory mechanisms occur in *both* heads—but in different ways.

#### 1. Inhibition of actin binding

Part of the actin-binding interface of the blocked head is obstructed by interaction with the free head, as shown previously [1] (interaction BF; red ellipse in Fig. 7C). In the free head, it is clear that actin-binding is also sterically blocked, through interaction of its actin-binding loop with the tail (interaction TF1; Fig. 5B; green ellipse in Fig. 7C, D), and by interference of the bundle of three tail segments (projecting up from TT1) with the free head actin-binding interface. Thus, each head would be prevented from binding to actin by distinct steric blocking mechanisms, one involving the tail and one the other head.

#### 2. Inhibition of ATPase activity

Similarly, ATPase activity of 10S myosin appears to be inhibited not only in the free, but also the blocked head, by stabilization of the converter domain of each, preventing phosphate release (cf. [52]). In the free head, the converter is immobilized by binding to the blocked head [1] (interaction BF; Fig. 5A; blue ellipse in Fig. 7A). In the blocked head, the converter may also be immobilized—in this case by binding to segment 2 of the tail as it wraps around the head (interaction TB3, Fig. 5A; red ellipse, Fig. 7A, B).

Inhibition of ATPase activity in both heads of 10S myosin would help explain a previously puzzling observation. Release of ATP hydrolysis products is ten times slower in 10S myosin than in HMM (0.0002 S^−1^ and 0.003 S^−1^ respectively) [31]. If the free head is inhibited in both (by binding to the blocked head), blocked head product release must be more inhibited in 10S myosin than in HMM. This could be explained by interaction of tail segment 2 with the blocked head converter in 10S myosin but not HMM, where it is missing. As mentioned previously, tail wrapping around the heads may also stabilize the basic head-head interaction of the IHM [1], enhancing the inhibitory effect of the two heads on each other in the 10S molecule.

We conclude that phosphate release and actin-binding are both inhibited in both heads, which can explain why 10S myosin is completely switched off [31], making it so well suited to its energy-conserving storage function in cells. The folded tail plays a critical role in this inhibition.

In thick filaments [23], both actin binding and ATPase are also sterically inhibited in both heads, again in different ways. Blocked-head actin binding is inhibited by interacting with the free head, as with 10S myosin, while the free-head is inhibited by orientation of its actin-binding face towards the thick filament backbone, away from actin. Both converters are also inhibited [23], impeding ATPase activity: in the free head by binding to the blocked head, as in 10S myosin; in the blocked head by interacting with S2 from the neighboring crown, as discussed above [23]. The greater number of interactions found in 10S myosin than in thick filaments suggests that the IHM of 10S myosin is more stable than in thick filaments, consistent with the greater inhibition observed in molecules [21, 31].

**Figure 7.**
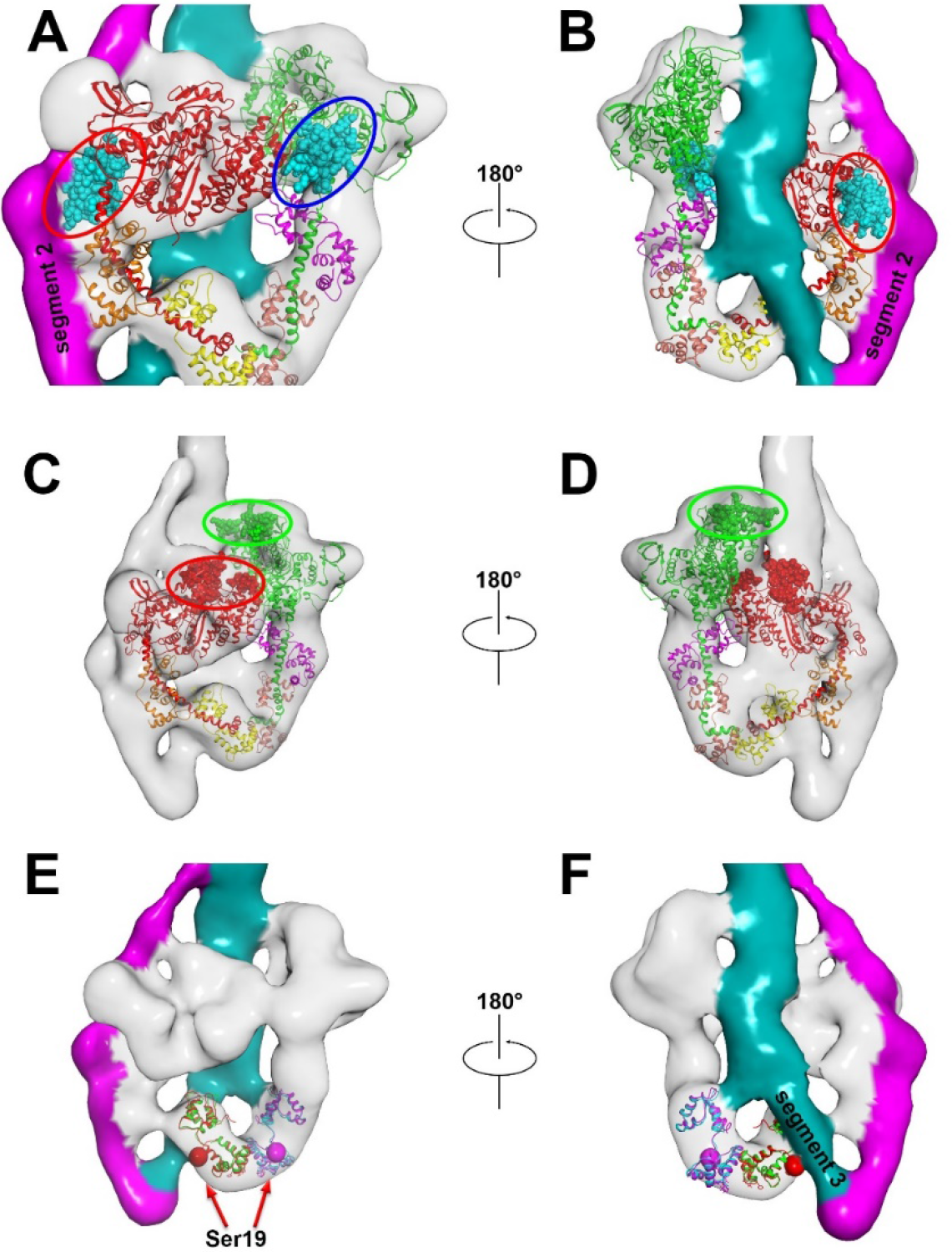
Inhibition mechanism of phosphate release and actin binding in the two heads. (A) Phosphate release in the blocked head is inhibited by interaction between its converter domain (cyan within red ellipse) and segment 2, and in the free head by interaction between its converter domain (cyan within blue ellipse) and the blocked head. (B) Back view, rotated 180° from (A). (C) and (D) Front and back views show actin-binding loop (red ellipse) in the blocked head sterically blocked by the free head, and actin-binding loop (green ellipse) in the free head blocked by interacting with the tail. (E) and (F) Front and back views showing steric blocking of Ser19 (red sphere) by segment 3 in the blocked head RLC, but not in the free head RLC (magenta sphere; see Movie 4).

### The central role of the tail in activation of 10S myosin

Smooth and nonmuscle 10S myosins are activated *in vitro* by phosphorylation of their RLCs. Under physiological ionic conditions this causes conversion to the 6S (unfolded) conformation [10, 15], which can form filaments, hydrolyze ATP and bind to actin. Our reconstruction provides insight into the mechanisms of these processes. Biophysical data suggest that phosphorylation of 10S myosin is a sequential process in which one RLC is phosphorylated more readily than the other, and it has been proposed that this might result from an asymmetric arrangement of the myosin heads [53, 54] (others have suggested that phosphorylation is random, but the starting myosin in those studies may not have been in the 10S conformation [55, 56]). The reconstruction reveals not only asymmetry of the heads (as previously described), but also dramatically different environments of their phosphorylation sites, which would readily account for a sequential phosphorylation process. Ser19 in the free head RLC is completely accessible to MLCK and would be easily phosphorylated. In the blocked head, Ser19 is sterically hindered by segment 3 of the tail (interaction TB5; Figs. 7E, F, Movie 4), which would hamper access of MLCK. Thus, the free head RLCs would be phosphorylated first, and blocked head RLCs later [53, 54, 57, 58]. A similar difference in accessibility of MLCK to the phosphorylatable serines is observed in tarantula filaments due to differing steric environments in the filament [41].

Biophysical measurements show that both RLCs must be phosphorylated for complete transition of the 10S to the 6S conformation [57, 58], and for significant activation of ATPase activity [53]. This suggests that phosphorylation of the blocked head RLC (occurring second) must ultimately determine unfolding, and that singly phosphorylated molecules would remain mostly folded and relatively inactive. The immediate proximity of Ser19 on the blocked head RLC to interaction TB5—which we suggested earlier is the critical interaction for creating the folded state (Fig. 7E, F)—would explain how phosphorylation of this serine could be the key step that causes tail unfolding. This proximity suggests a direct (rather than allosteric) effect of phosphorylation on the binding of segment 3: decreased positive charge on the RLC would weaken its interaction with the negatively charged region of segment 3, favoring unfolding of the tail [35]. Extension of RLC helix A upon phosphorylation [46], or changes in ELC-RLC interaction affecting the conformation of the light chain domain [48, 59], might also contribute to the loss of interaction with segment 3. Importantly, heavy meromyosin, which lacks segments 2 and 3, requires phosphorylation of only one of its two heads for activation [60]: in this case TB5 does not exist, and phosphorylation of a single head is sufficient to break the weaker head-head interaction in HMM. The requirement for phosphorylation of both heads in 10S myosin (one of which is sterically inhibited) for full activation may be another mechanism by which the cell conserves energy in the quiescent state. If any MLCK molecules in the cell are Ca^2+^-insensitive (e.g. through proteolysis or other damage), resulting RLC phosphorylation will occur preferentially on the free head, but this will not activate the molecule. Only when intracellular Ca^2+^ levels rise will there be sufficient activated MLCK to phosphorylate the blocked head RLC as well, with concomitant unfolding of the 10S structure.

Based on the above observations and previous work, we propose the following mechanism for regulating 10S myosin activity in cells. **1.** At low intracellular Ca^2+^, both heads are dephosphorylated, and the molecule is in the 10S conformation, in which ATPase activity and actin binding are inhibited by the mechanisms discussed earlier. **2.** With increased Ca^2+^, MLCK is activated, and first phosphorylates the unobstructed RLC of the free head. This may alter the conformation of the light chain domain of this head, weakening interaction of the free head with the blocked head and the tail, while retaining the overall folded conformation and relatively low ATPase activity of the molecule [9, 48, 53, 57–59]. **3.** The less accessible blocked head RLC is then phosphorylated. This could result from the weakening of the IHM just mentioned, or from “breathing” of the intact 10S structure, which transiently exposes blocked head Ser19 (we assume that the interactions we have described are relatively weak and in equilibrium between interacting and non-interacting states). There is also evidence that the RLCs may interact with each other and with the flexible initial portion of subfragment 2, stabilizing the conformation of this region of the IHM [23, 28, 50]. Phosphorylation of the free head RLC may disrupt these interactions, leading to increased local flexibility and greater accessibility of Ser19 on the blocked head RLC. **4.** Phosphorylation of the blocked head phosphorylation domain weakens or abolishes its interaction (TB5) with the negative charge cluster on segment 3 due to decreased charge attraction. **5.** With breakage of TB5 (the key interaction required for the folded state), the binding of segments 1 and 2 to the heads is destabilized, allowing the tail to become fully extended. **6.** Loss of head-tail interactions during tail unfolding may further destabilize previously weakened head-head contacts, promoting separation of the heads. **7.** The extended tails now interact to form filaments, actin-binding sites on both heads are exposed, and inhibitory interactions of the converter domains are lifted, leading to full actin-activated ATPase activity and filament sliding.

## Materials and Methods

### Electron microscopy

Smooth muscle myosin II from turkey gizzard was purified according to [61]. Smooth muscle myosin has similar structural and functional properties to nonmuscle myosin II and is a convenient source for our experiments on the 10S structure [10]. Myosin molecules in 0.15 M NaAc, 1 mM EGTA, 2.5 mM MgCl_2_, 0.5 mM ATP, 10 mM MOPS, pH 7.5 were negatively stained at 10 nM concentration using 1% (w/v) uranyl acetate [33]. Negative staining is a simple and powerful technique for obtaining 3D structural data at moderate resolution. While it does not provide the high resolution possible with cryo-EM, it is more than adequate for revealing structure and interactions at the domain level and was therefore our method of choice (see [33, 34]); the detail observed in our reconstruction was sufficient to provide multiple new insights into the mechanism of inhibition in the 10S molecule. Carbon films were pretreated with UV light to optimize stain spreading [2, 62]. Molecules were lightly crosslinked in solution at room temperature for 1 min with 0.1% glutaraldehyde before staining [2, 35]. This was designed to minimize distortion caused by binding to the carbon surface and drying of the stain, and does not significantly alter the structure of the molecules at the resolution of our studies [2, 35]. 200 micrographs were recorded under low dose conditions on a Philips CM120 electron microscope at 120 kV with a 2K × 2K CCD camera (F224HD, TVIPS GmbH) at a pixel size of 3.7 Å (nominal 45,000 magnification). These micrographs were used for 3D reconstruction and refinement. 100 pairs of micrographs of tilted (50°) and untilted (0°) grids were collected at 5.4 Å/pixel (nominal magnification 37,000) for random conical tilt reconstruction.

### Image processing

#### Initial model from random conical tilt

No complete 3D model is available for 10S myosin II to use as a reference for 3D reconstruction. To exclude model bias, we implemented the random conical tilt method [36] to create a starting model, using SPIDER [63]. 13,675 pairs of particles were interactively selected from the 100 tilted and untilted micrographs using the program JWEB in SPIDER. 2D classification was performed for the untilted particles. Four classes of particle with similar views in the class averages were chosen for further analysis. For each class, the Euler angle of each particle from the tilt images was calculated based on the tilt angle and the azimuthal angle from 2D classification. Back projection was carried out to compute the 3D reconstruction. Finally, the four class averages were rotated to make the tail parallel to the y-axis. The obtained azimuthal angles were used to modify the Euler angles and to merge particles from the chosen four classes, and the model was built using the merged data set.

#### 2D classification and 3D reconstruction

The folded tail of 10S myosin protruding beyond the heads is flexible, and also rotates (within a 60° range) about its junction with the heads [9]. To simplify analysis, only particles with an unbent tail pointing straight up from the head-head junction were chosen for image processing. Particles with dimensions 150 × 150 pixels, including both heads and a small portion of the protruding tail, were manually selected using EMAN [64], and then low-pass filtered to 20 Å. All subsequent image processing was carried out with RELION [40]. Four rounds of 2D classification were performed. Bad or low-resolution particles with smeared densities were removed after each round. The remaining 15833 particles were subjected to 3D classification into 6 classes. Classes 1, 2 and 5 showed variable appearances of the tail and heads. Classes 3 and 4 had similar 3D reconstructions and were combined for refinement. The resolution of the refined 3D reconstruction was estimated to be 25 Å (Fig. S2). Class 6 molecules showed less detail than classes 3 and 4, and were excluded from the final reconstruction

#### Atomic fitting

The atomic model of the interacting heads of chicken smooth muscle myosin (PDB 1I84) [1] was docked into the 3D reconstruction using rigid-body fitting in CHIMERA [65] (Fig. 4). Molecular dynamics flexible fitting was not attempted owing to the low resolution. Densities outside of the heads were attributed to the three segments of the tail. According to 3D classification (Fig. 6), S2 in the 3D reconstruction was quite flexible. We therefore chose not to fit an atomic model of S2 to the reconstruction.

## Acknowledgments

We thank Drs. Edward Korn and Xiong Liu for *Dictyostelium* myosin and Sanford Bernstein and Floyd Sarsoza for insect flight muscle myosin. We thank the Electron Microscopy Facility and its staff at UMass Medical School for instrumentation and assistance. This work was supported by NIH grants AR072036, AR067279 and HL139883 (RC) and HL075030 and HL111696 (MI). MI is an awardee of University of Texas STARS PLUS Award. The content is solely the responsibility of the authors and does not necessarily represent the official views of the National Institutes of Health.

## SUPPLEMENTARY FIGURES

**Figure S1.**
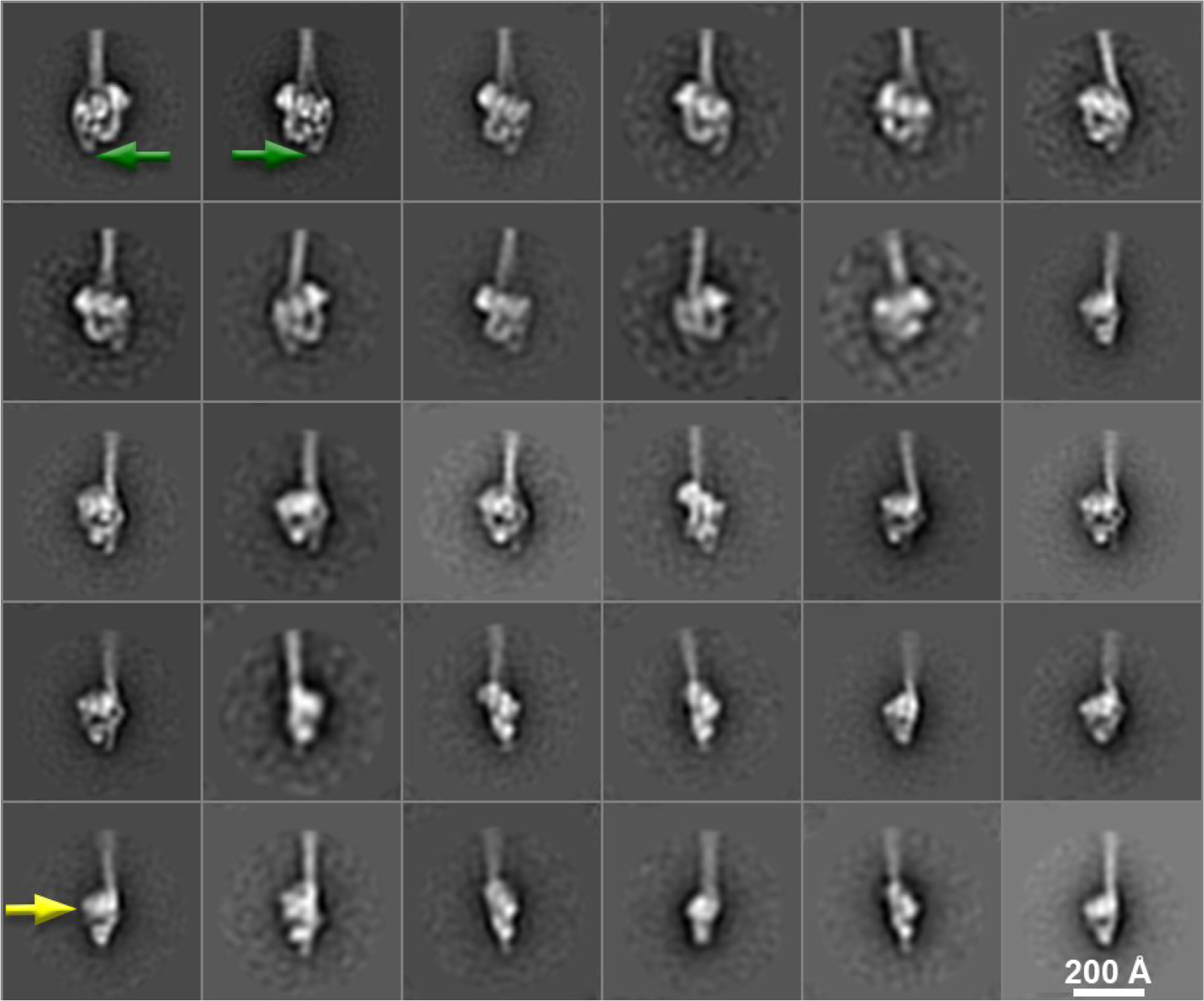
2D class averages of negatively stained 10S myosin molecules. The appearances of the different class averages indicate that the monomers tend to lie on the grid with the tail parallel to the grid surface, but with different rotations about its long axis. Mirrored particles indicated by green arrows represent face views of molecules lying face-up or face-down [2]; arrows point specifically to hinge 2 (Fig. 1). Yellow arrow points to a side view.

**Figure S2.**
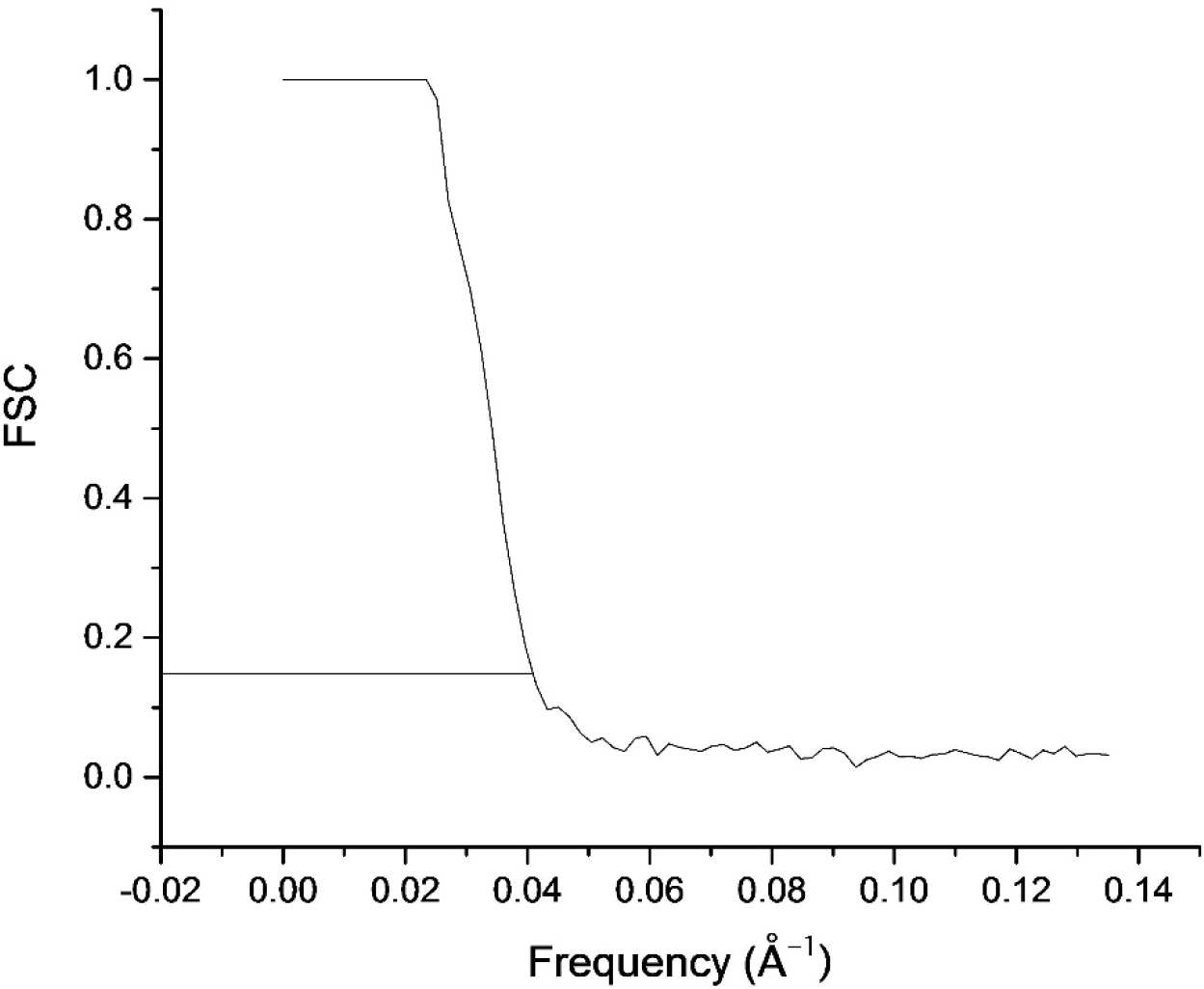
Gold-standard Fourier shell correlation (FSC) curve to determine resolution.

**Figure S3.**
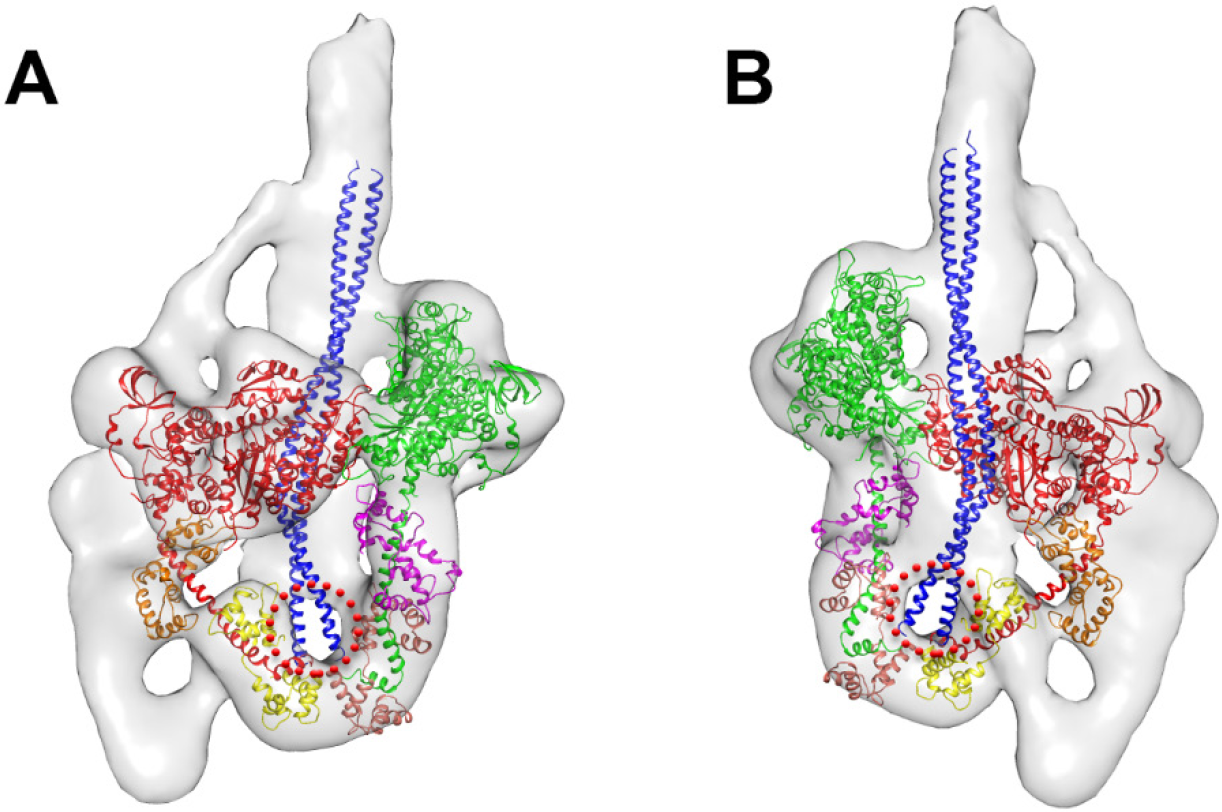
Location of initial portion of subfragment 2. Front and back views (A and B, respectively) show the expected course of subfragment 2 (blue ribbon, based on ref. 21), starting at the junction between the blocked and free head RLCs. The red dotted circle indicates an absence of density in the reconstruction for the initial part of S2, probably due to mobility of this part of the coiled-coil (see text and Fig. 3E).

**Figure S4.**
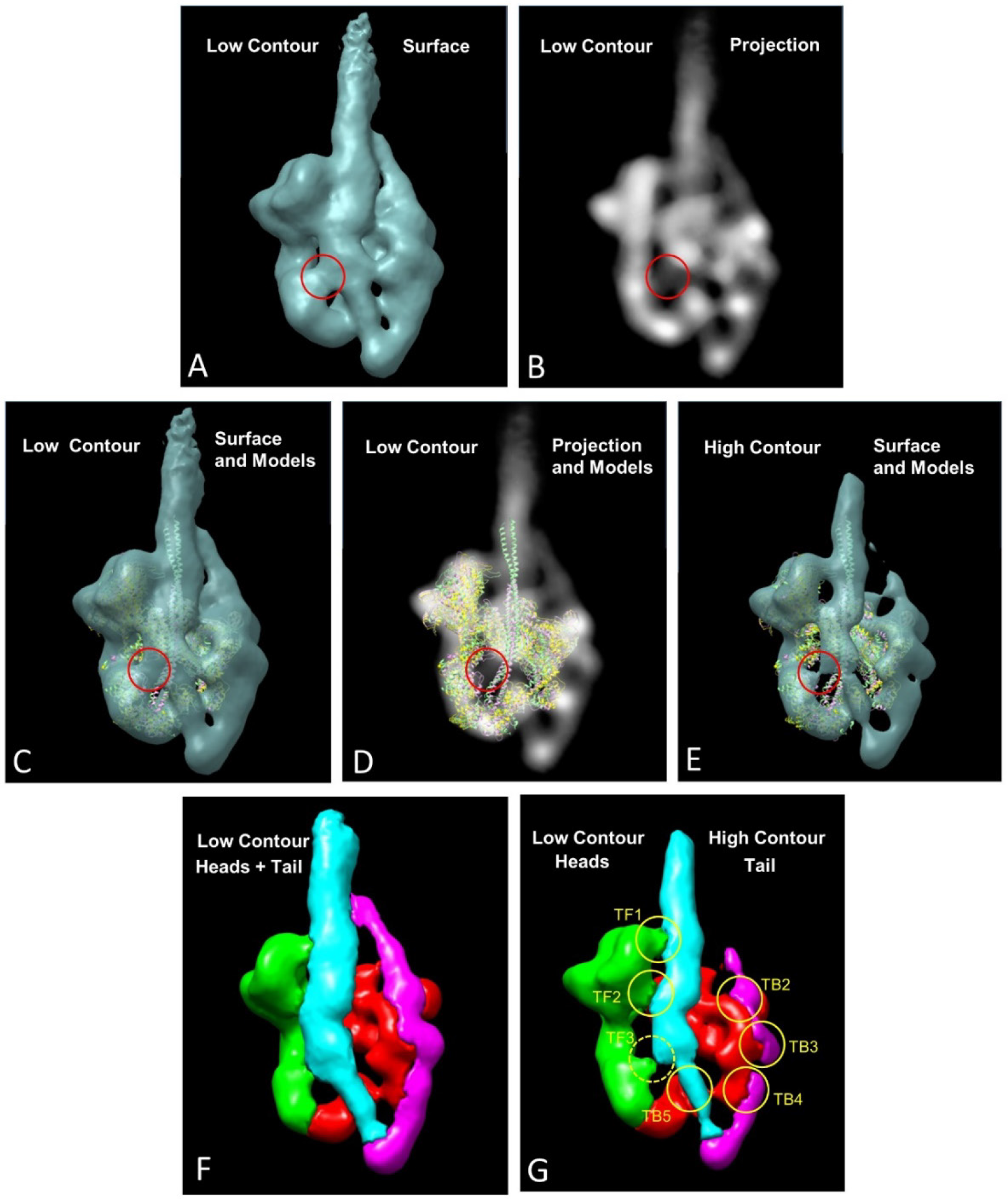
Comparison of low and high contour cutoff reconstructions reveals the stronger and weaker parts of the density (rear views). (A, B) Surface and projection (density) views. Apparent mass in low contour cutoff (circled interaction TF3 in (A)) is seen to have very little density in (B). (C-E) show TF3 in low (C) and high (E) contour cutoff surface views and in projection (D), with different variants of the IHM/S2 model fitted as in Fig. S3. Note disappearance of TF3 at high contour, and absence of protein density to fill the TF3 volume (circles). Note also difference in subfragment 2 density (bottom right of circle) between (C) and (E). Compare with Fig. S3. (F, G) With high contour cutoff, flexible regions of tail disappear (G), while contacts with the heads remain, suggesting that these are real interactions. The exception is TF3, which disappears.

**Figure S5.**
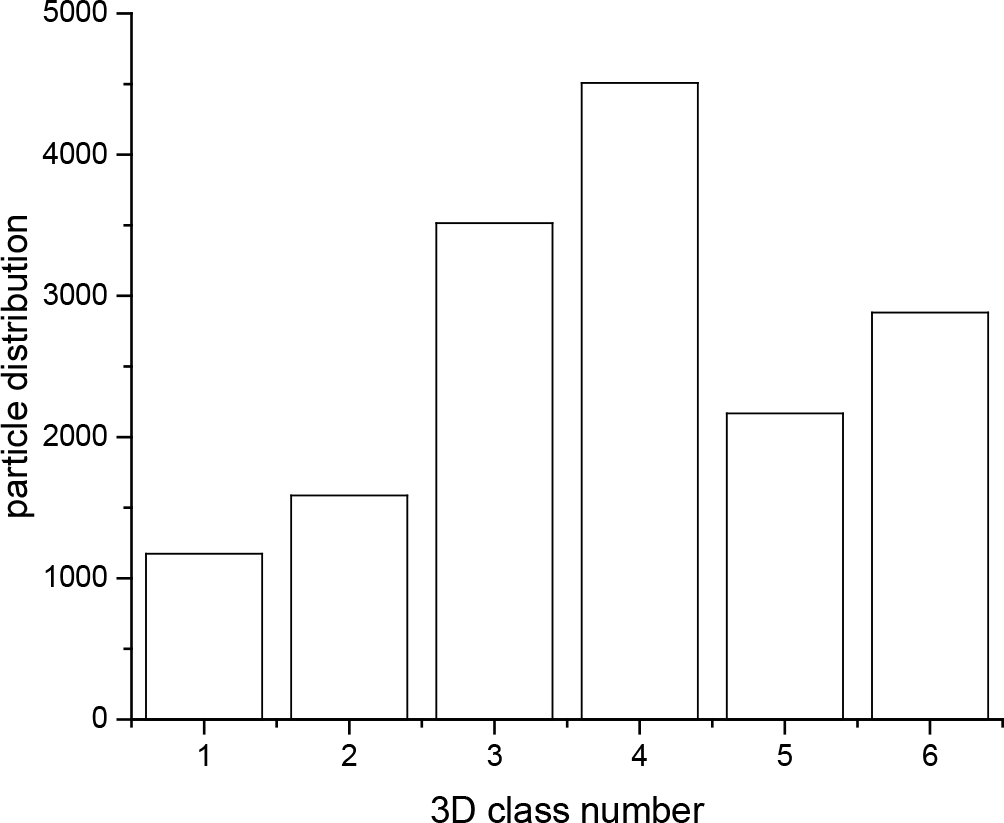
Particle distribution of the 6 classes after 3D classification. The corresponding 3D reconstructions are shown in Fig. 6.

**Figure S6.**
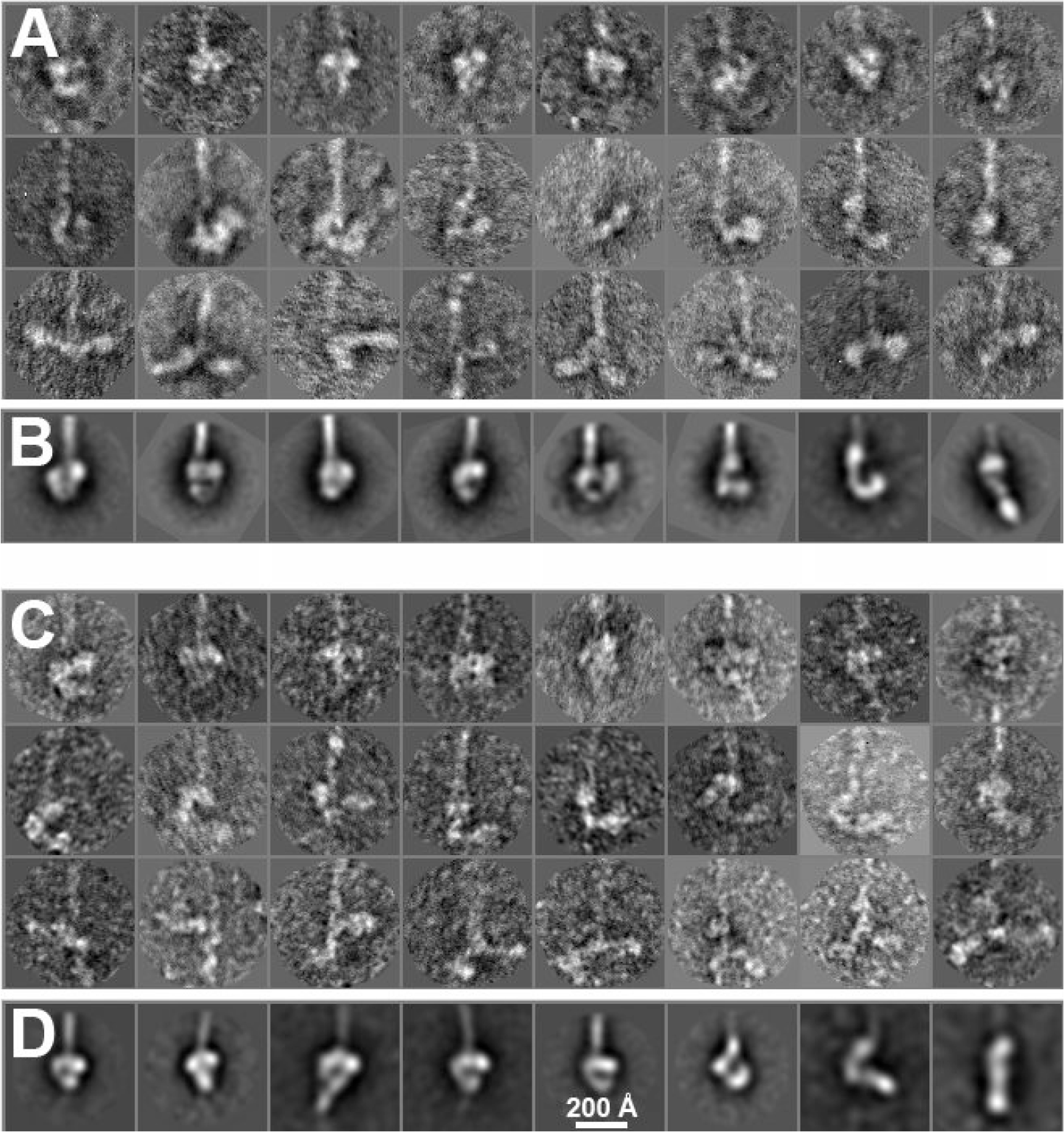
Negatively stained insect indirect flight muscle (A,B) and *Dictyostelium* myosin II (C,D; unpublished data from [26]). The top rows of (A) and (C) show molecules with a compact head arrangement in the IHM conformation. The second rows show dissociation of the free head from the blocked head, with the latter usually attached to subfragment 2. The free head exhibits gradual dissociation from the blocked head from left to right. The third rows show an open structure with no intramolecular interactions. (B) and (D) are 2D class averages of flight muscle and *Dictyostelium* myosin, respectively, computed to show the variability of head organization. The IHM is clearly demonstrated in the left images, while some averages on the right show the free head separated from the blocked head. Interaction of segments 2 and 3 with the heads in smooth muscle 10S myosin considerably stabilizes the IHM. The above myosins, in which these interactions are weak or absent, serve to illustrate this point. Smooth muscle myosin clearly shows the second hinge point in 2D averages, a characteristic marker of the three-segment folded structure (green arrows, Fig. S1). The absence of this feature in many insect flight myosin molecules suggests that segments 2 and 3 are more weakly bound to the heads. Correspondingly, the free head tends to detach from the blocked head and the blocked head also becomes more mobile, as shown in (C) above. In *Dictyostelium*, segment 3 continues past the two heads rather than making a second bend [26], and segment 2 does not appear to make interactions around the blocked head [26]. Correspondingly, free and blocked heads again were frequently dissociated from each other and the tail.

**Figure S7.**
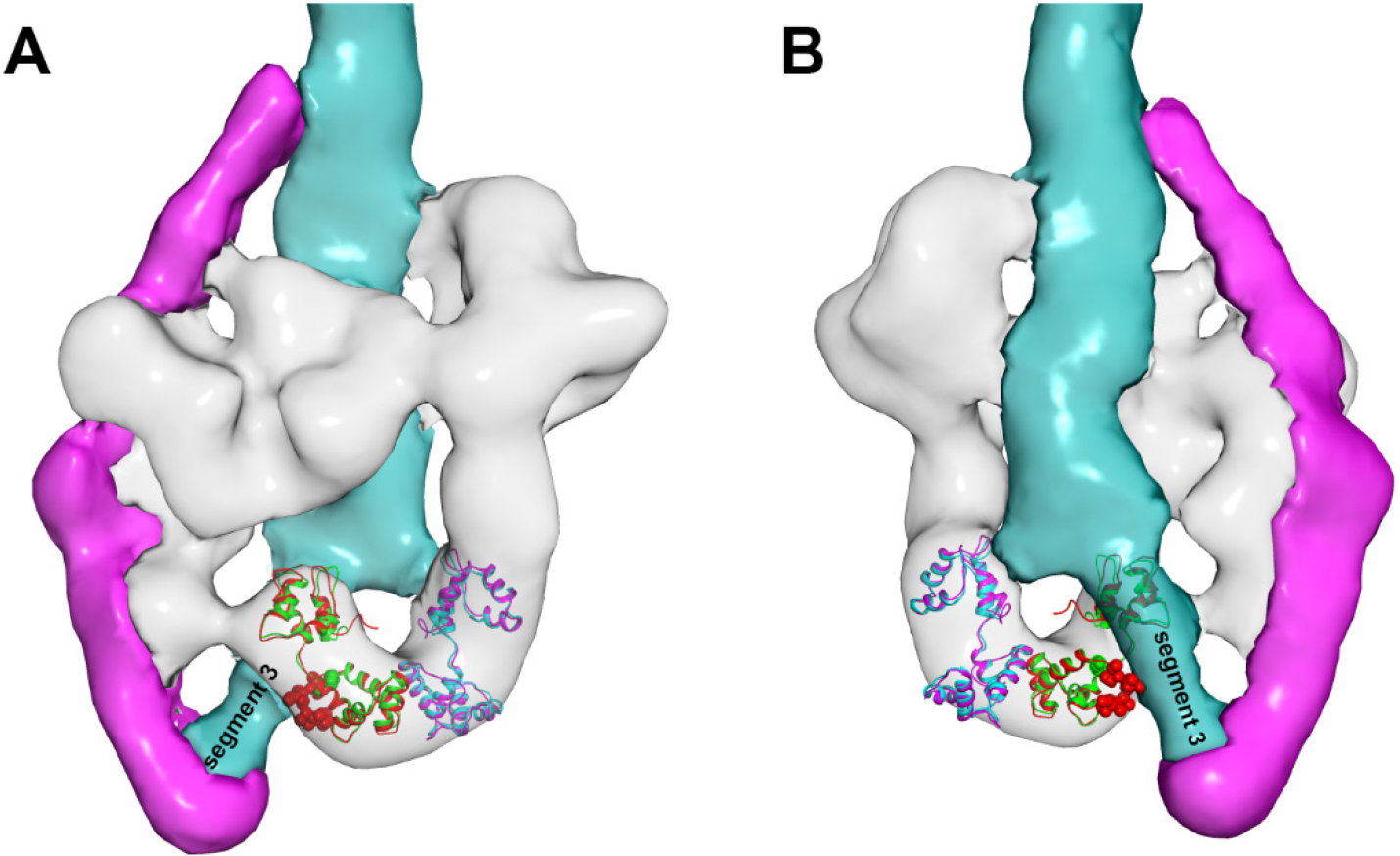
The RLC N-terminal interacts with tail segment 3 in the 10S molecule. The N-terminal 24 residues of the smooth muscle RLC are absent in crystal structures due to mobility (see text). To estimate their location, we docked the scallop RLC (3JTD; red, which has more residues present) onto the smooth muscle RLC (green), so that the location of the additional residues could be visualized. Residues 1-11 (red spheres, equivalent to residues 14-24 in smooth muscle) suggest that the RLC N-terminal directly interacts with segment 3. A and B are front and back views.

## SUPPLEMENTARY MOVIES

**Movie 1. 3D reconstruction of 10S smooth muscle myosin rotated about its long axis (see Fig. 3)**. Blocked head is red; free head, green; tail segments 1 and 3, cyan; segment 2, magenta.

**Movie 2. Atomic fitting of PDB-1I84 into the 3D reconstruction of 10S molecules (see Fig. 4).** The densities attributed to the tail can be explicitly distinguished after atomic fitting. The path of the tail wrapping around the heads can also be easily traced.

**Movie 3. Tail segment 3 contacts the RLC of the blocked head (see Fig. 5).** Segment 3 is seen to closely approach Cys108 (cyan sphere) in the C-terminal lobe of the blocked head RLC.

**Movie 4. Tail segment 3 sterically block Ser19 on the blocked head (see Fig. 7).** Segment 2, magenta; segment 3, cyan. Blocked head Ser19, red; free head Ser19, magenta. Note: The blocked and free head RLCs of 1i84 are colored green and blue respectively. However, these chains end at residue 25, owing to disorder of the N-terminus. To estimate the position of Ser19 in each case, we superimposed the scallop RLC (red and magenta) on the smooth muscle RLC, and have marked residue 19 in each case.

## References

1. Wendt, T., Taylor, D., Trybus, K.M., and Taylor, K. (2001). Three-dimensional image reconstruction of dephosphorylated smooth muscle heavy meromyosin reveals asymmetry in the interaction between myosin heads and placement of subfragment 2. Proceedings of the National Academy of Sciences of the United States of America 98, 4361–4366.

2. Jung, H.S., Komatsu, S., Ikebe, M., and Craig, R. (2008). Head-head and head-tail interaction: a general mechanism for switching off myosin II activity in cells. Mol Biol Cell 19, 3234–3242.

3. Yang, S., Woodhead, J.L., Zhao, F.Q., Sulbaran, G., and Craig, R. (2016). An approach to improve the resolution of helical filaments with a large axial rise and flexible subunits. J Struct Biol 193, 45–54.

4. Geeves, M.A., and Holmes, K.C. (1999). Structural mechanism of muscle contraction. Annual review of biochemistry 68, 687–728.

5. Vicente-Manzanares, M., Ma, X., Adelstein, R.S., and Horwitz, A.R. (2009). Non-muscle myosin II takes centre stage in cell adhesion and migration. Nat Rev Mol Cell Biol 10, 778–790.

6. Tajsharghi, H., and Oldfors, A. (2013). Myosinopathies: pathology and mechanisms. Acta Neuropathol 125, 3–18.

7. Moore, J.R., Leinwand, L., and Warshaw, D.M. (2012). Understanding cardiomyopathy phenotypes based on the functional impact of mutations in the myosin motor. Circulation research 111, 375–385.

8. Suzuki, H., Onishi, H., Takahashi, K., and Watanabe, S. (1978). Structure and function of chicken gizzard myosin. Journal of biochemistry 84, 1529–1542.

9. Burgess, S.A., Yu, S., Walker, M.L., Hawkins, R.J., Chalovich, J.M., and Knight, P.J. (2007). Structures of smooth muscle myosin and heavy meromyosin in the folded, shutdown state. Journal of molecular biology 372, 1165–1178.

10. Craig, R., Smith, R., and Kendrick-Jones, J. (1983). Light-chain phosphorylation controls the conformation of vertebrate non-muscle and smooth muscle myosin molecules. Nature 302, 436–439.

11. Suzuki, H., Kamata, T., Onishi, H., and Watanabe, S. (1982). Adenosine triphosphate-induced reversible change in the conformation of chicken gizzard myosin and heavy meromyosin. J Biochem 91, 1699–1705.

12. Trybus, K.M., Huiatt, T.W., and Lowey, S. (1982). A bent monomeric conformation of myosin from smooth muscle. Proc Natl Acad Sci U S A 79, 6151–6155.

13. Cross, R.A., Cross, K.E., and Sobieszek, A. (1986). ATP-linked monomer-polymer equilibrium of smooth muscle myosin: the free folded monomer traps ADP.Pi. The EMBO journal 5, 2637–2641.

14. Cross, R.A. (1988). What is 10S myosin for? Journal of muscle research and cell motility 9, 108–110.

15. Trybus, K.M., and Lowey, S. (1984). Conformational states of smooth muscle myosin. Effects of light chain phosphorylation and ionic strength. The Journal of biological chemistry 259, 8564–8571.

16. Sellers, J.R. (1991). Regulation of cytoplasmic and smooth muscle myosin. Current opinion in cell biology 3, 98–104.

17. Seow, C.Y. (2005). Myosin filament assembly in an ever-changing myofilament lattice of smooth muscle. American journal of physiology. Cell physiology 289, C1363–1368.

18. Milton, D.L., Schneck, A.N., Ziech, D.A., Ba, M., Facemyer, K.C., Halayko, A.J., Baker, J.E., Gerthoffer, W.T., and Cremo, C.R. (2011). Direct evidence for functional smooth muscle myosin II in the 10S self-inhibited monomeric conformation in airway smooth muscle cells. Proceedings of the National Academy of Sciences of the United States of America 108, 1421–1426.

19. Xu, J.Q., Gillis, J.M., and Craig, R. (1997). Polymerization of myosin on activation of rat anococcygeus smooth muscle. Journal of muscle research and cell motility 18, 381–393.

20. Takahashi, T., Fukukawa, C., Naraoka, C., Katoh, T., and Yazawa, M. (1999). Conformations of vertebrate striated muscle myosin monomers in equilibrium with filaments. Journal of biochemistry 126, 34–40.

21. Ankrett, R.J., Rowe, A.J., Cross, R.A., Kendrick-Jones, J., and Bagshaw, C.R. (1991). A folded (10 S) conformer of myosin from a striated muscle and its implications for regulation of ATPase activity. J Mol Biol 217, 323–335.

22. Katoh, T., Konishi, K., and Yazawa, M. (1998). Skeletal muscle myosin monomer in equilibrium with filaments forms a folded conformation. The Journal of biological chemistry 273, 11436–11439.

23. Woodhead, J.L., Zhao, F.Q., Craig, R., Egelman, E.H., Alamo, L., and Padron, R. (2005). Atomic model of a myosin filament in the relaxed state. Nature 436, 1195–1199.

24. Stewart, M.A., Franks-Skiba, K., Chen, S., and Cooke, R. (2010). Myosin ATP turnover rate is a mechanism involved in thermogenesis in resting skeletal muscle fibers. Proc Natl Acad Sci U S A 107, 430–435.

25. Jung, H.S., Burgess, S.A., Billington, N., Colegrave, M., Patel, H., Chalovich, J.M., Chantler, P.D., and Knight, P.J. (2008). Conservation of the regulated structure of folded myosin 2 in species separated by at least 600 million years of independent evolution. Proc Natl Acad Sci U S A 105, 6022–6026.

26. Lee, K.H., Sulbaran, G., Yang, S., Mun, J.Y., Alamo, L., Pinto, A., Sato, O., Ikebe, M., Liu, X., Korn, E.D., et al. (2018). Interacting-heads motif has been conserved as a mechanism of myosin II inhibition since before the origin of animals. Proc Natl Acad Sci U S A 115, E1991–E2000.

27. Pinto, A., Sanchez, F., Alamo, L., and Padron, R. (2012). The myosin interacting-heads motif is present in the relaxed thick filament of the striated muscle of scorpion. Journal of structural biology 180, 469–478.

28. Woodhead, J.L., Zhao, F.Q., and Craig, R. (2013). Structural basis of the relaxed state of a Ca2+-regulated myosin filament and its evolutionary implications. Proc Natl Acad Sci U S A 110, 8561–8566.

29. Zhao, F.Q., Craig, R., and Woodhead, J.L. (2009). Head-head interaction characterizes the relaxed state of Limulus muscle myosin filaments. Journal of molecular biology 385, 423–431.

30. Zoghbi, M.E., Woodhead, J.L., Moss, R.L., and Craig, R. (2008). Three-dimensional structure of vertebrate cardiac muscle myosin filaments. Proc Natl Acad Sci U S A 105, 2386–2390.

31. Cross, R.A., Jackson, A.P., Citi, S., Kendrick-Jones, J., and Bagshaw, C.R. (1988). Active site trapping of nucleotide by smooth and non-muscle myosins. Journal of molecular biology 203, 173–181.

32. Liu, J., Wendt, T., Taylor, D., and Taylor, K. (2003). Refined model of the 10S conformation of smooth muscle myosin by cryo-electron microscopy 3D image reconstruction. Journal of molecular biology 329, 963–972.

33. Takizawa, Y., Binshtein, E., Erwin, A.L., Pyburn, T.M., Mittendorf, K.F., and Ohi, M.D. (2017). While the revolution will not be crystallized, biochemistry reigns supreme. Protein Sci 26, 69–81.

34. Burgess, S.A., Walker, M.L., Thirumurugan, K., Trinick, J., and Knight, P.J. (2004). Use of negative stain and single-particle image processing to explore dynamic properties of flexible macromolecules. Journal of structural biology 147, 247–258.

35. Jung, H.S., Billington, N., Thirumurugan, K., Salzameda, B., Cremo, C.R., Chalovich, J.M., Chantler, P.D., and Knight, P.J. (2011). Role of the tail in the regulated state of myosin 2. J Mol Biol 408, 863–878.

36. Radermacher, M., Wagenknecht, T., Verschoor, A., and Frank, J. (1987). Three-dimensional reconstruction from a single-exposure, random conical tilt series applied to the 50S ribosomal subunit of Escherichia coli. Journal of microscopy 146, 113–136.

37. Fabiola, F., and Chapman, M.S. (2005). Fitting of high-resolution structures into electron microscopy reconstruction images. Structure 13, 389–400.

38. Brown, J.H., Yang, Y., Reshetnikova, L., Gourinath, S., Suveges, D., Kardos, J., Hobor, F., Reutzel, R., Nyitray, L., and Cohen, C. (2008). An unstable head-rod junction may promote folding into the compact off-state conformation of regulated myosins. Journal of molecular biology 375, 1434–1443.

39. Spudich, J.A. (2015). The myosin mesa and a possible unifying hypothesis for the molecular basis of human hypertrophic cardiomyopathy. Biochem Soc Trans 43, 64–72.

40. Scheres, S.H. (2012). RELION: implementation of a Bayesian approach to cryo-EM structure determination. Journal of structural biology 180, 519–530.

41. Brito, R., Alamo, L., Lundberg, U., Guerrero, J.R., Pinto, A., Sulbaran, G., Gawinowicz, M.A., Craig, R., and Padron, R. (2011). A molecular model of phosphorylation-based activation and potentiation of tarantula muscle thick filaments. Journal of molecular biology 414, 44–61.

42. Trybus, K.M., Freyzon, Y., Faust, L.Z., and Sweeney, H.L. (1997). Spare the rod, spoil the regulation: necessity for a myosin rod. Proceedings of the National Academy of Sciences of the United States of America 94, 48–52.

43. Olney, J.J., Sellers, J.R., and Cremo, C.R. (1996). Structure and function of the 10 S conformation of smooth muscle myosin. The Journal of biological chemistry 271, 20375–20384.

44. Trybus, K.M., and Lowey, S. (1988). The regulatory light chain is required for folding of smooth muscle myosin. The Journal of biological chemistry 263, 16485–16492.

45. Trybus, K.M., and Chatman, T.A. (1993). Chimeric regulatory light chains as probes of smooth muscle myosin function. The Journal of biological chemistry 268, 4412–4419.

46. Espinoza-Fonseca, L.M., Kast, D., and Thomas, D.D. (2008). Thermodynamic and structural basis of phosphorylation-induced disorder-to-order transition in the regulatory light chain of smooth muscle myosin. Journal of the American Chemical Society 130, 12208–12209.

47. Ikebe, M., Ikebe, R., Kamisoyama, H., Reardon, S., Schwonek, J.P., Sanders, C.R., 2nd, and Matsuura, M. (1994). Function of the NH2-terminal domain of the regulatory light chain on the regulation of smooth muscle myosin. The Journal of biological chemistry 269, 28173–28180.

48. Taylor, K.A., Feig, M., Brooks, C.L., 3rd, Fagnant, P.M., Lowey, S., and Trybus, K.M. (2014). Role of the essential light chain in the activation of smooth muscle myosin by regulatory light chain phosphorylation. J Struct Biol 185, 375–382.

49. Nag, S., Trivedi, D.V., Sarkar, S.S., Adhikari, A.S., Sunitha, M.S., Sutton, S., Ruppel, K.M., and Spudich, J.A. (2017). The myosin mesa and the basis of hypercontractility caused by hypertrophic cardiomyopathy mutations. Nat Struct Mol Biol 24, 525–533.

50. Alamo, L., Wriggers, W., Pinto, A., Bartoli, F., Salazar, L., Zhao, F.Q., Craig, R., and Padron, R. (2008). Three-dimensional reconstruction of tarantula myosin filaments suggests how phosphorylation may regulate myosin activity. Journal of molecular biology 384, 780–797.

51. Alamo, L., Qi, D., Wriggers, W., Pinto, A., Zhu, J., Bilbao, A., Gillilan, R.E., Hu, S., and Padron, R. (2016). Conserved Intramolecular Interactions Maintain Myosin Interacting-Heads Motifs Explaining Tarantula Muscle Super-Relaxed State Structural Basis. J Mol Biol 428, 1142–1164.

52. Yount, R.G., Lawson, D., and Rayment, I. (1995). Is myosin a “back door” enzyme? Biophysical journal 68, 44S–47S; discussion 47S-49S.

53. Persechini, A., and Hartshorne, D.J. (1981). Phosphorylation of smooth muscle myosin: evidence for cooperativity between the myosin heads. Science 213, 1383–1385.

54. Persechini, A., and Hartshorne, D.J. (1983). Ordered phosphorylation of the two 20 000 molecular weight light chains of smooth muscle myosin. Biochemistry 22, 470–476.

55. Trybus, K.M., and Lowey, S. (1985). Mechanism of smooth muscle myosin phosphorylation. The Journal of biological chemistry 260, 15988–15995.

56. Wagner, P.D., Vu, N.D., and George, J.N. (1985). Random phosphorylation of the two heads of thymus myosin and the independent stimulation of their actin-activated ATPases. The Journal of biological chemistry 260, 8084–8089.

57. Ikebe, M., Hinkins, S., and Hartshorne, D.J. (1983). Correlation of intrinsic fluorescence and conformation of smooth muscle myosin. The Journal of biological chemistry 258, 14770–14773.

58. Ikebe, M., Hinkins, S., and Hartshorne, D.J. (1983). Correlation of enzymatic properties and conformation of smooth muscle myosin. Biochemistry 22, 4580–4587.

59. Houdusse, A., and Cohen, C. (1996). Structure of the regulatory domain of scallop myosin at 2 A resolution: implications for regulation. Structure 4, 21–32.

60. Walcott, S., Fagnant, P.M., Trybus, K.M., and Warshaw, D.M. (2009). Smooth muscle heavy meromyosin phosphorylated on one of its two heads supports force and motion. The Journal of biological chemistry 284, 18244–18251.

61. Ikebe, M., and Hartshorne, D.J. (1985). Effects of Ca2+ on the conformation and enzymatic activity of smooth muscle myosin. The Journal of biological chemistry 260, 13146–13153.

62. Burgess, S.A., Walker, M.L., Thirumurugan, K., Trinick, J., and Knight, P.J. (2004). Use of negative stain and single-particle image processing to explore dynamic properties of flexible macromolecules. J. Struct. Biol 147, 247–258.

63. Frank, J. (2006). Three-dimensional electron microscopy of macromolecular assemblies: visualization of biological molecules in their native state, 2nd Edition, (Oxford; New York: Oxford University Press).

64. Tang, G., Peng, L., Baldwin, P.R., Mann, D.S., Jiang, W., Rees, I., and Ludtke, S.J. (2007). EMAN2: an extensible image processing suite for electron microscopy. J Struct Biol 157, 38–46.

65. Pettersen, E.F., Goddard, T.D., Huang, C.C., Couch, G.S., Greenblatt, D.M., Meng, E.C., and Ferrin, T.E. (2004). UCSF Chimera--a visualization system for exploratory research and analysis. Journal of computational chemistry 25, 1605–1612.

